# Bacteria have numerous phage-plasmid families with conserved phage and variable plasmid gene repertoires

**DOI:** 10.1101/2020.11.09.375378

**Authors:** Eugen Pfeifer, Jorge A. Moura de Sousa, Marie Touchon, Eduardo P.C. Rocha

## Abstract

Plasmids and temperate phages are mobile genetic elements driving bacterial evolution. They are usually regarded as very distinct. However, some elements, termed phage-plasmids, are known to be both plasmids and phages, *e*.*g*. P1, N15 or SSU5. The number, distribution, relatedness and characteristics of these phage-plasmids are poorly known. Here, we screened for these elements among ca. 14000 phages and plasmids and identified 780 phage-plasmids across very diverse bacterial phyla. We grouped 92% of them by similarity of gene repertoires to define 8 families and 18 other broader communities of elements. The existence of these large groups suggests that phage-plasmids are ancient. Their gene repertoires are large, the average element is larger than an average phage or plasmid, and they include slightly more homologs to phages than to plasmids. We analyzed the pangenomes and the genetic organization of each group of phage-plasmids and found the key phage genes to be conserved and co-localized within families, whereas genes with homologs in plasmids are much more variable and include most accessory genes. Phage-plasmids are a sizeable fraction of all phages and plasmids and could have key roles in bridging the genetic divide between phages and other mobile genetic elements.

## INTRODUCTION

The evolution of Bacteria to novel challenges is facilitated by their ability to acquire genes by horizontal gene transfer, a process typically mediated by self-mobilizable genetic elements. These elements can be distinguished based on the mechanism of horizontal transmission between cells and of vertical transmission within cellular lineages. Horizontal transmission usually takes place either by conjugation or within virions (1). The latter may follow diverse mechanisms: either the temperate phage integrates the novel genome as a prophage or it transduces bacterial DNA following one of several distinct mechanisms (2, 3). Most genes in prophages are silent, but some may be expressed and confer novel phenotypes to the lysogen (lysogenic conversion). Vertical transmission of mobile genetic elements (MGEs) takes place by autonomous replication of plasmids or by their integration in the chromosome. The textbook view is that conjugative elements tend to be plasmids (4), whereas temperate phages, such as lambda, tend to integrate the chromosome as prophages (5). Yet, it is now known that the majority of conjugative MGEs integrates the chromosome as ICEs (6).

It has also been known for decades that some functional temperate phages integrate the genome as extra-chromosomal plasmids that replicate in line with the cell cycle (7–9). These prophages are thus also plasmids. Here, we shall follow Ravin et al., (10) and call them phage-plasmids (P-P). P-Ps have functions that are typically associated with plasmids to replicate and segregate at each cell division. For this, they require an initiator of replication (11) (such as a replicase). Some small high-copy number plasmids rely only on passive diffusion for segregation between daughter cells, but model P-Ps are large replicons and are therefore expected to encode partition systems (12). Because P-Ps are temperate phages, they can infect bacteria, produce virions, lyse the host and infect other host cells. Hence, they need to encode many of the typical functions of temperate phages: regulation of lysogeny, lysis, DNA packaging, and virion structure. Contrary to chromosomal prophages, P-Ps do not need to encode recombinases for site-specific recombination with the chromosome (typically integrases). However, they may encode recombinases to resolve dimers, as many plasmids (13), or to alternate between an integrative and a plasmid state (8). Finally, known P-Ps encode accessory functions often identified in large MGEs, such as defense (7) or toxin-antitoxin systems (7, 14). Some elements strongly resembling experimentally demonstrated P-Ps have genes encoding for virulence factors (15), antibiotic resistance (16), or the capsule (15).

The first reported P-Ps - P1 and N15 - were isolated over fifty years ago (14, 17) and became established model systems in the field of molecular biology. P1 is widely used as a strong general transducer (18), because its headful packaging system (Pacase) occasionally incorporates host DNA into the virion (19). P1 also encodes the site-specific Cre-recombinase to resolve head-to-tail multimers (7), which has become a versatile tool in genetic engineering (20, 21). N15 has a linear dsDNA genome with covalently closed ends produced by a protelomerase (TelN) (22), and is a model system to study the formation, resolution and diversity of linear replicons in Bacteria (23). A few P-Ps closely related to P1 and N15 have been reported (9, 24), but their numbers and diversity are poorly known. Other P-Ps have been described in enterobacteria, mycobacteria, *Vibrio, Bacillus* and *Clostridiales* (9, 25–28). A noteworthy case is the phage SSU5, that was isolated from a *Salmonella enterica* strain (29) and is a promising auxiliary component of phage cocktails (30). A comparative analysis based on a few strains revealed that this phage is related to plasmids encoding proteins homologous to phage sequences that are found in distantly related hosts such as *Escherichia, Klebsiella* and *Yersinia* (31). P1, N15 and SSU5 represent only a few examples of potential P-Ps. Several plasmids were reported to have genes homologous to phages and some phages to have genes homologs to plasmids (e.g. pHCM2, pECOH89, RHEph10 and SJ46 (32–35)). Whether these correspond to P-Ps is usually unknown.

The abundance of P-Ps, their relatedness, and their gene contents are poorly known. Two studies have identified elements with nucleotide sequence similarity to P1, SSU5 (9) and N15 (36). Here, we aim at identifying and characterizing P-Ps using more sensitive analyses of protein homology to assess their distribution across Bacteria. The identification of distant homologs allowed to search systematically for phage functions in known plasmids and for plasmid functions in known phages, resulting in the identification of a large number of putative P-Ps. This finding spurred three questions. Can these elements be grouped in meaningful families? Are P-Ps more like phages or more like plasmids? How do gene repertoires vary across different groups? To answer these questions, we clustered P-Ps by similarity of gene repertoires, defined P-P families, characterized their functions, and used them to study the frequency of phage-like functions relative to plasmid-like functions in P-Ps. Our results show that P-Ps are very diverse in terms of the size, function and organization of gene repertoires.

## MATERIAL AND METHODS

### Data and data processing

The complete genomes of 11827 plasmids (with accompanying bacterial genomes) and 2502 phages were retrieved from NCBI non-redundant RefSeq database (37) (ftp://ftp.ncbi.nlm.nih.gov/genomes/refseq/, last accessed in May 2019). The information on the virus family for the phages was taken from the GenBank file under the ORGANISM description (50 phages were unassigned in the file). The replicons were assigned to a bacterial host species using the GenBank file (under ORGANISM) for plasmids and the virus-host database (https://www.genome.jp/virushostdb/) for phages. Additionally, we downloaded 12230 phage genomes from the main section of GenBank that passed a quality filter and were absent from RefSeq. This database has many highly similar phages and was only used to search for homologs of representative P-Ps. It was retrieved from the Virus database of NCBI (https://www.ncbi.nlm.nih.gov/labs/virus/vssi/#/) (38) (last accessed in August 2020). All analysis and visualization were conducted in the R environment (39), if not otherwise stated.

### Annotation of protein sequences

The functional annotation of protein sequences was done using HMMER v3.b2 (40) searches with default parameters to the PFAM (41) (version 32.0, September 2018, https://pfam.xfam.org/), TIGRFAM (42) (version 15.0, September 2014, http://tigrfams.jcvi.org/cgi-bin/index.cgi), eggNOG (43) (bactNOG and Viruses only) (version 5.0 November 2018, http://eggnog5.embl.de/#/app/home) and pVOG (44) (version 1, first May 2017, http://dmk-brain.ecn.uiowa.edu/pVOGs/home.html#) databases (downloaded in May 2019). We used the ‘--cut_ga’ option when searching for homology to profiles of the PFAM and TIGFRAM databases to restrict the hits to those with reliable scores. If not otherwise stated positive hits were assigned using the same criteria as used by MacSyFinder (45) (profile coverage >= 50%, idomain_evalue <= 10^−3^).

### Database of phage-specific HMM profiles

The phage-specific profiles were carefully chosen from pVOG, PFAM and TIGRFAM databases. The pVOG database has phage-specific HMMs with information on their viral quotient (VQ) (44). The VQ ranges from 0 to 1 and indicates the specificity of the pVOG to viruses. A value of VQ close to 1 means that the profile matches almost only virus genomes, whereas a value close to 0 means most matches are from cellular genomes (44). To complement the pVOG database with profiles that are curated manually, we combined it with the PFAM and TIGRFAM databases. First, a reciprocal profile-profile comparison was conducted between all 9518 pVOGs and phage specific PFAMs (n=366) and TIGRFAMs (n=71) (phage-specific PFAM and TIGRFAM profiles were taken from Phage_Finder (46)) using HHsearch (47) (included in HHsuite 2.0.9) with a significance p-value threshold of 10^−5^. Only bidirectional hits were considered. The 437 PFAM/TIGRFAM profiles clustered with 711 pVOGs into 260 groups (based on Louvain community detection (48), singletons excluded). These profiles are designated as group 1 profiles in the training of the random forest model (suppl. table 1) (see below). Second, pVOG profiles built from alignments with at least 15 sequences and with a VQ higher than 0.75 (n=1435). These profiles are designated as group 2 profiles in the training of the random forest model (suppl. table 2) (see below). The 2583 group 1 and 2 phage-specific profiles were classed in six categories a) structure, b) lysis, c) packaging, maturation/assembly and DNA injection, d) recombination, regulation and DNA metabolism e) unknown and f) others.

### Identification of phage-plasmids (P-P)

To identify P-Ps we screened known phages for plasmid-associated functions and known plasmids for phage-associated functions. We excluded ssDNA phages (*Ino*- and *Microviridae*), elements smaller than 10 kb (smallest dsDNA phage in RefSeq) and larger than 300 kb (to avoid megaplasmids that might have been integrated by temperate phages).

We searched phages for plasmid-associated genes using HMMs specific for plasmid replication (38 profiles from (49)) and plasmid partition systems (9 profiles from (49) and 48 from databases, suppl. table 3). Genes associated with conjugation, *i*.*e*. the mating pair formation apparatus and the relaxase, were searched using CONJscan (50). This resulted in the identification of 122 phages that contained plasmid features (suppl. table 4).

The plasmid database was screened by random forest prediction models to identify P-Ps. Ideally, one would have learned the models on P-Ps as positives and other plasmids as negatives. However, the number of elements experimentally demonstrated to be P-Ps is too small. Hence, we made an approximation and built models that were trained to distinguish plasmids predicted to lack phage functions (negatives) and known phages (positives). The training datasets included 2000 randomly chosen phages (positives) and 2000 randomly chosen plasmids lacking prophage fragments (negatives). The latter are those were PHASTER (51) (ran with default parameters) could not identify intact, questionable or incomplete prophages.

The plasmids were searched for hits to the categorized phage-specific profiles (described in “database of phage-specific HMM profiles”, suppl. table 1 and 2). We computed 16 different fractions (per replicon: the number of hits in a category was divided by the overall number of proteins) from these results: six functional categories of phage-specific group 1 (n=6), same for group 2 HMM profiles (n=6), pVOG HMMs, phage-PFAM and phage-specific TIGRFAM profiles and fractions of proteins lacking hits. In addition, the number of proteins per replicon was considered (as a control). These 17 features were used for training and evaluation, which was conducted using the ranger (52) package in R. The parameters used to train the models were set to: 10 000 trees, “mtry” = √(feature number) = 4 (number of variables to possibly split at each node was set to default), “splitrule” to “extratrees” and the computation of the variable “importance” mode is based on “permutation” (Fig S1). The type of forest (“treetype”) was chosen to be “regression” to assign a probability - phage probability score (PSC) - that ranges between 0 and 1. A score close to “0” indicates a plasmid lacking phage genes and a score close to “1” indicates that the plasmid has a high probability of also being a phage. To achieve a higher accuracy, we repeated this approach 10 times to build 10 models.

In each round, we kept a test dataset (independent from the train dataset), consisting of 4950 plasmids (lacking prophage fragments as predicted by PHASTER) and 497 phages. The out of the box error (O.O.B.) was about 1.3 % ± 0.1 % (Fig. S1A). Subsequently, the 10 test datasets were used to validate the 10 models with the pROC package (53) in R. In this evaluation, each model was applied on a test dataset independent from its own training dataset. The area under the curve (AUC) based on the receiver operating characteristics - true positive rate (sensitivity) vs false positive rate (specificity) - was ∼0.99 (Fig S1B).

The 10 models were used to class plasmids that were predicted by PHASTER to contain fragments of prophages. Each plasmid was analyzed in the light of each of the 10 models, leading to 10 PSCs values that were averaged per plasmid. We found 566 potential P-Ps with a mean PSC > 0.5 (suppl. table 4). This list was complemented with putative P-Ps from the literature (see main text).

### Sequence similarity network of phage-plasmids: Construction, clustering, curation

We searched for significant similarity (e-value <10^−4^, identity ≥ 35%, coverage ≥ 50%) among all pairs of P-P proteins using MMseqs2 (version 9-d36de) (54). The best bi-directional hits (BBH) between pairs of elements were used to calculate the weighted gene repertoire relatedness (wGRR) (49, 55):

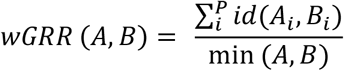

Where A_i_ and B_i_ are the *i*th BBH pair of *P* total pairs, the number of genes from the smaller P-P is min(A,B), and the identity between the BBH pair is id(A_i_,B_i_). The wGRR varies between 0 (no BBH) and 1 (all genes in an element have an identical BBH in the other).

The wGRR scores were used to compute a sequence similarity network of the putative P-Ps (suppl. table 5). Genome pairs with a wGRR <= 0.05 were discarded to reduce the signal’s noise. The communities of P-Ps in the network were detected using the Louvain algorithm (48) with the NetworkToolbox (56) (R package). The default gamma parameter (γ=1) was increased to γ=1.9 to split some large communities (for a study on the variation on this parameter see Fig S2). The clustering resulted in 26 communities with three or more P-Ps (Fig. 3), 7 doubletons and 47 singletons (Fig. S3). Communities, that are made of members found only in the plasmid database, were screened for related phages from the non-redundant GenBank phage database to identify cases of high wGRR similarity. Phages with a wGRR score of at least 0.1 to a P-P were considered as related (suppl. table 6). We then defined families (and eventually subfamilies) of P-Ps within communities with more than two members. Of note, the communities are assigned by the Louvain algorithm, whereas families are subsets of communities where were weakly related P-Ps were removed. Communities that were too small or too diverse (e.g. PiSa, Actinophage A) or lacking key functions (e.g. cp32) were not curated (Fig. S4). The separation of P-Ps within a community into different subfamilies (or their exclusion from the family) is based on the analysis of the persistent genome: members of a family have at least 10% of persistent genes (see pangenome detection below, Fig S8-S13). Overall, this process of curation led to the identification of one P-P superfamily, 10 families and 4 subfamilies (suppl. tables 7 and 8).

### Typing phage-plasmids in terms of plasmid incompatibility and virus taxonomy

We used PlasmidFinder 2.0.1 (57) with default parameters to class the incompatibility types of P-Ps (suppl. table 4). The virus taxonomies of P-Ps identified from the phage database were retrieved from the GenBank file under the ORGANISM entry. P-Ps identified from the plasmid database were classed using the hits to pVOG profiles (n=9518) as features in a random forest model using the ranger (52) package in R. We used. Training and evaluation were done as for the prediction of phage-like features in plasmids (see “Identification of P-Ps” and text S1). We trained 10 models using 2000 randomly chosen phages (positives, known taxonomy) and 2000 randomly chosen plasmids with a mean PSC < 0.1 (negatives, taxonomy was set to plasmid-like). Each model gave probability scores for all possible taxonomies and only the one with the highest probability was considered. The computed out of the box (O.O.B.) prediction error was 3.0% ± 0.1 %. The evaluation of the 10 models was done by 10 data sets, each with 500 randomly chosen phages and 1000 randomly chosen plasmids (not in the training dataset). The correct assignment rate was 98.4% (Fig. S5A). For the elements tested by at least three out of ten models, the probability average was 98.8% with a standard deviation of 0.2% (Fig. S5B). We classed P-Ps when the average values of the probability minus the standard deviation were higher than 0.5 (suppl. table 4). In a few cases the class of the highest probability assignment differed among the 10 models. In these cases, we chose the taxonomy with the highest frequency. If the P-P was classed “plasmid-like”, the virus taxonomy was left unassigned (Fig. S5C).

### Calculation of the phage-plasmid quotient (PPQ)

MMseqs2 (version 9-d36de) (54) was used to calculate the similarity between all proteins of phages, plasmids and P-Ps (e-value <10^−4^, identity >= 35%, coverage ≥ 50%). The BBHs were extracted and used to compute the phage-plasmid quotient (PPQ) per protein sequence. BBHs with plasmids with PSC > 0.1 were removed to avoid searching for similarity to degenerated P-Ps (or potential P-Ps lacking many known phage genes). Genes lacking homologs in phages or plasmids were excluded. The PPQ scores were computed according to the following equation (see text S2):

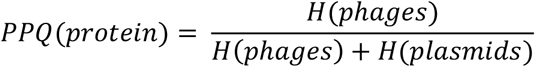

Where *H(phages)* is the number of BBH between P-Ps and phages normalized to the size of the phage database and *H(plasmids)* is the same quantity relative to the plasmid database. The PPQ represents the preponderance of phage hits (relative to plasmids). It is calculated per protein sequence and varies between 0 (mostly plasmid hits) and 1 (mostly phage hits). We computed a PPQ per P-P (gPPQ) by making the average of the PPQs for the P-P (elements with less than 10 protein sequences with a PPQ score were excluded, Fig. S7, suppl. tables 4). The gPPQ varies between 0 (mostly like a plasmid) and 1 (mostly like a phage).

### Computation and visualization of pangenomes

Pangenomes were calculated using PPanGGOLiN (58) version 1.0.1 with default parameters except for the AB subfamily g2 where the parameter for the max degree smoothing was set to 2 (default = 10) because this group contains only five members. This program uses MMseqs2 (54) to cluster proteins with more than 80% amino acid identity and 80% coverage. PPanGGOLiN then calculates the presence/absence (P/A) matrix of the gene families and performs a partitioning of the families into persistent (present in most P-Ps of a family), shell (present in an intermediate number of P-Ps), and cloud genomes (present in few P-Ps). These matrices were used to curate the P-P communities into families or subfamilies (Fig S8-S13). Indexed pangenomes graphs (Figures S15-S19) were visualized and inspected using Gephi (59) (as recommended by (58)) and the igraph (60) package in R. Additional information on the gene families of the pangenomes are given in supplemental tables 9 to 19.

The pangenome graphs were colored using the average sequence similarity of the BBH across the P-P (sub) family to produce similarity pangenome graphs (self-hits excluded) (e.g. Fig. S9C) or colored using the average PPQ values of each gene family to produce PPQ pangenome graphs (e.g. Fig 5A). This allows to identify the variability of gene repertoires in the light of the function, relatedness within P-P family and frequency of the gene family in the pangenomes.

## RESULTS AND DISCUSSION

### Many phage-plasmids in databases

We screened the RefSeq database (14 329 phages and plasmids) for putative P-Ps, excluding ssDNA phages, plasmids smaller than 10 kb (the smallest tailed dsDNA phage is 11 kb; NC_002515) and larger than 300 kb (may be megaplasmids or chromids with prophages). This resulted in a set of 11 284 phages and plasmids. In the absence of published methods to identify P-Ps, we developed an approach to identify phage core functions in known plasmids and another approach to identify plasmid core functions in known phages. We assumed that such elements are good P-P candidates. We searched 2383 known phages for genes involved in plasmid replication, partition and conjugation. We detected 122 putative P-Ps (Fig. 1A), including most of the already reported elements (e.g. P1, N15 and SSU5, suppl. table 4). Some known elements that are absent from RefSeq such as P7, D6 and pMCR-1-P3 (9, 61) were also correctly identified by our methods in a complementary analysis of GenBank replicons absent from RefSeq. Yet, for consistence and to avoid redundancy, we only present the data concerning RefSeq.

**Figure 1:**
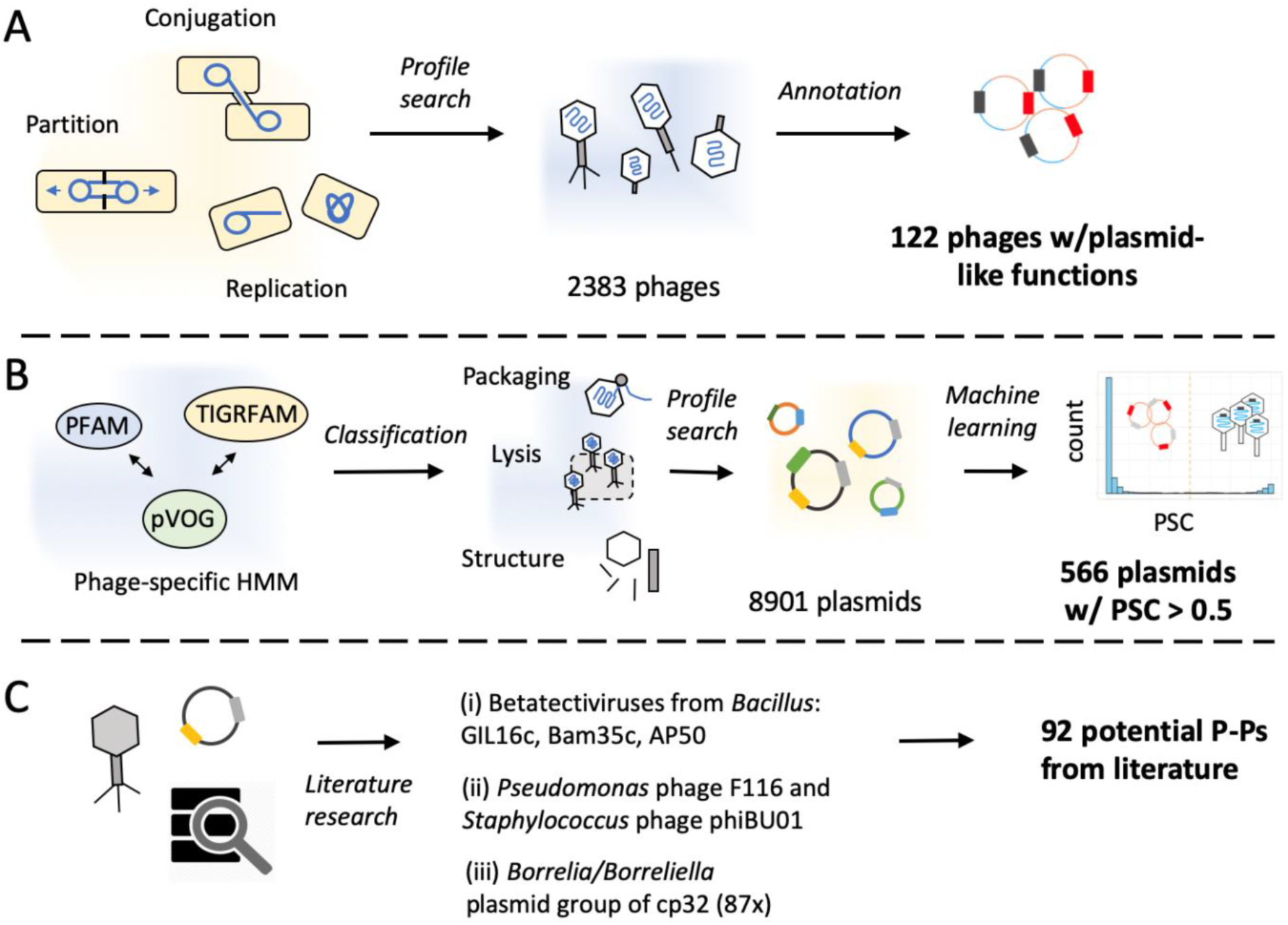
Methods to screen databases for P-Ps. **A**. 2383 phages were annotated using protein profiles specific of plasmids (conjugation, replication and partition), resulting in 122 putative P-Ps (suppl. table 4). **B**. Carefully selected phage protein profiles were classified into distinct phage-specific functions such as structure components, packaging/maturation, lysis, etc. Their hits were used as features to train a machine learning method to distinguish plasmids lacking any kind of prophage from phages. We then used 10 random forest models to screen a plasmid database of 8901 plasmids yielding in 566 putative P-Ps (phage probability score (PSC) > 0.5) (suppl. table 4). **C**. The screen for P-Ps was complemented by searching the literature for other potential P-Ps.

We used a machine learning approach to identify phage-associated traits in 8901 known plasmids. For this, we made a database of 2583 phage-associated protein profiles and used them to find such genes in plasmids (Fig. 1B). We then trained random forest models to distinguish phages from plasmids lacking any kind of prophage regions (see Methods). These models revealed high sensitivity, high specificity and a low error rate (1.3%±0.01%, Figure S1AB). Replicons with phage probability score (PSC) > 0.5 were regarded as putative P-Ps. We found 566 such putative P-Ps among known plasmids (Figure S1E).

We searched the literature for P-Ps missed in our screen. These rare cases could be classed in three groups (Fig1 C, sup Text 1): (i) The three linear dsDNA Betatectiviruses GIL16c, Bam35c and AP50 (27, 62), lack recognizable plasmid functions. Yet, the linear plasmids (pBClin15 from *B. cereus* or pBMBLin15 from *B. thuringiensis* (27, 62)) are closely related and have identifiable phage-related functions. Hence, P-Ps in this group could be identified when scanning known plasmids, but not when scanning known phages. (ii) Two distinct phages with known extra-chromosomal replicative states - F116 from *Pseudomonas* and phiBU01from *Staphyloccocus* (8, 63) - lack recognizable plasmid related functions. Closely related elements were absent from the plasmid database. (iii) The cp32-like *Borellia/Borreliella* plasmids have been proposed to be P-Ps (64). These 87 elements lack recognizable key phage-related functions: head, tail or capsid proteins (PSC ranging from 0 to 0.38). A phage, phi-BB-1, found among cp-32 like elements is capable of forming virions and transduce DNA of a different cp32 (65). We thus assumed that these three groups of elements are known or putative P-Ps. Together with the P-Ps from the two screening approaches this resulted in a set of 780 P-Ps (suppl. table 4). Although we may have missed P-Ps, especially in poorly studied phyla, we can already conclude that P-Ps are numerous. They are 7.3% of the plasmids and 5.6% of the phages of RefSeq.

### Phage-plasmids are prevalent and have a bimodal size distribution

P-Ps occur in many bacterial species scattered across 81 host genera (suppl. table 4). They can be found in Firmicutes such as *Bacillus* and *Clostridium*, in Actinobacteria such as in *Mycobacterium*, and in alpha, beta and gamma Proteobacteria like enterobacteria, *Vibrio, Acinetobacter, Zymomonas*, and *Burkholderia* (Fig. 2A). More than 200 of the P-Ps are found in *Escherichia* and *Klebsiella* species, where they represent 7.3% of their ∼2900 phages and plasmids. This is consistent with a report where ∼7% of the *E. coli* strains had P1-like P-Ps (16). A large number of P-Ps was also found in mycobacterial phages. Although no P-P was detected among the few available mycobacterial plasmids (n=72), we found plasmid functions, mostly partition systems, in 6.5% of the 365 mycophages. These P-Ps belong to the huge cluster A of temperate actinophages (25), of which 20% lack an integration module (25, 66). The frequency of P-Ps among phages and plasmids is even higher in other less sampled clades. For example, P-Ps are a large fraction (up to 50%) of phages or plasmids of *Arsenophonus, Bacillus, Clostridia* and *Piscirickettsia*. Two of these genera, *Bacillus* and *Clostridia*, have many sequenced genomes, which means that high frequency of P-P is not an artifact due to small samples. We found more than one P-P element in 75 bacterial genomes of which most are in *Klebsiella, Piscirickettsia* and *Bacillus* (suppl. table 4).

**Figure 2:**
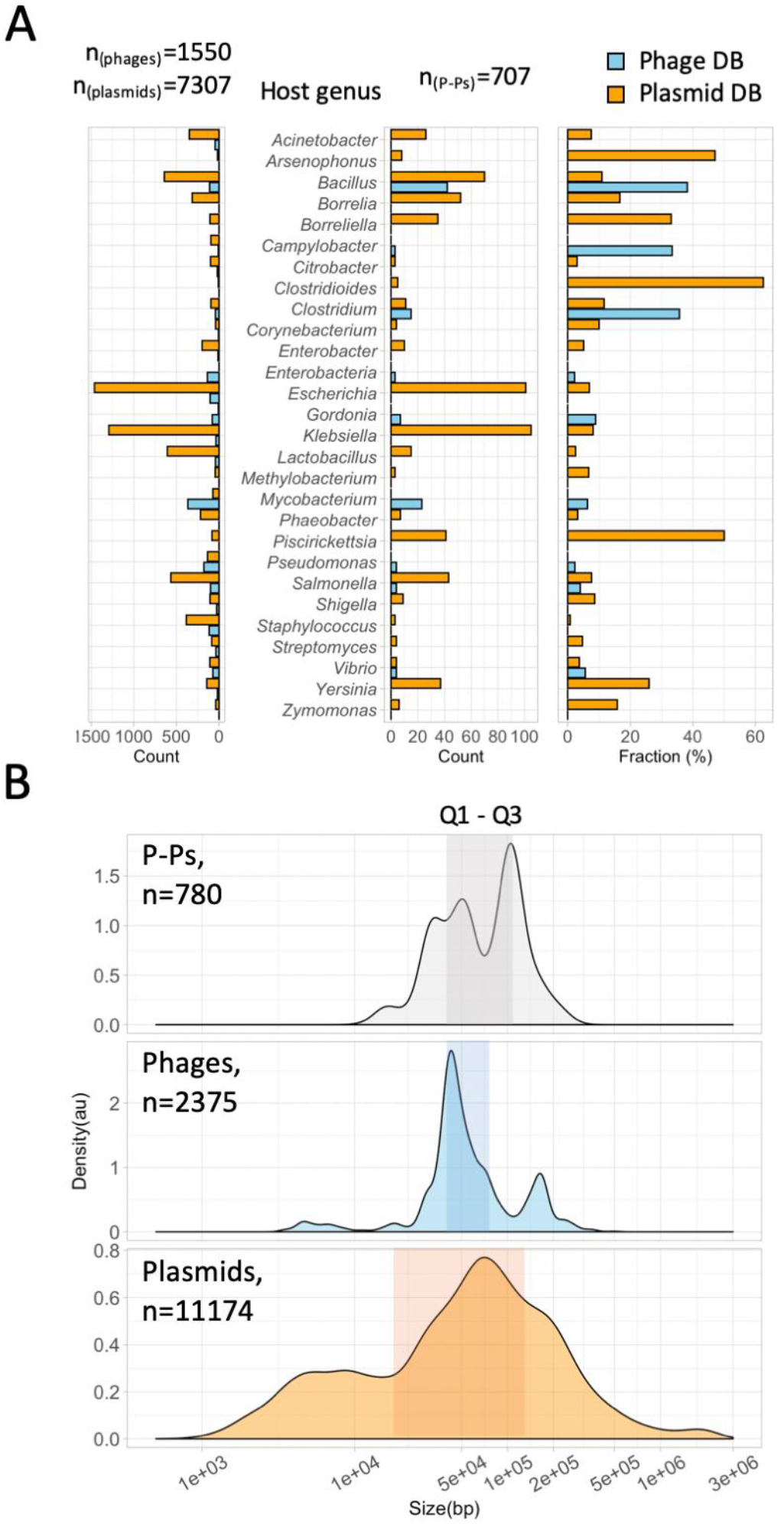
Host and size distribution of P-Ps. **A**. The frequency in bacterial genera of phages and plasmids (left) and P-Ps (center), for genera with at least three P-Ps (for the complete host distribution see suppl. table 4). The right panel shows the frequency of P-Ps per host genus normalized to the sizes of the databases of phages and plasmids. **B**. Density plots (mean normalized counts) of replicon sizes from P-Ps (grey), phages (blue) and plasmids (orange). The shaded boxes indicate the ranges of the first (Q1, 25%) and the third (Q3, 75%) quantiles representing 50% of the replicons.

The distribution of P-P genome sizes shows two interesting patterns (Fig. 2B). First, its bimodal with a broad peak around 50 kb and a sharper one around 100 kb. Their average size (median_P-Ps_= 67.8 kb) is larger than those of both plasmids (median_Plasmids_= 59.1 kb) and phages (median_Phages_= 48.5 kb). Presumably this is because P-Ps have to encode the key functions of both types of elements. Second, the quantiles of this distribution are intermediate from the ones of plasmids and phages. On average, the interquartile distance of P-P genome sizes (ΔQ1,Q3= 68.8 kb) is almost half that of plasmids (ΔQ1,Q3= 112.6 kb) and double that of phages (ΔQ1,Q3= 35.8 kb). It’s likely that contrary to plasmids, sudden changes in P-P size are restricted by the need to accommodate its genome in the virion.

### Composition and diversity of P-P families

We searched for homology across the gene repertoires of the 780 P-Ps by computing the weighted gene repertoire relatedness (wGRR), which integrates information on the presence of homologs and their sequence identity (see Methods, suppl table 5). It varies between zero (no homologs) and one (all genes from one element have an identical homolog in the other). Among pairs with moderate to high wGRR values (>0.05), many include comparisons of P-Ps from the phage and the plasmid databases (mean_wGRR_= 0.32 to be compared with an average of 0.38 for comparisons between P-Ps from the phage database), showing that the two sources of P-Ps have many homologous elements.

We clustered the wGRR matrix using the Louvain algorithm (48) and detected 26 communities with at least three P-Ps. The communities were named after representative members (e.g. P1, N15, SSU5) or the clade of the most frequent host (e.g. AB for *<u>A</u>cinetobacter <u>b</u>aumannii*, PiSa for *<u>Pi</u>scirickettsia <u>sa</u>lmonis*) (Fig. 3). Five communities showed high wGRR values within and between the communities and were classed into one supercommunity (named after SSU5). From the remaining 21 communities, three large (cp32, PiSa and pLP39) and five small ones (less than 10 members) are made of members that were identified only in the plasmid database (suppl. table 4). These putative P-Ps contained key phage functions (in agreement with their classification as phages by our random forest model). To search for known phages related to those ‘plasmid-only’ communities, we screened phages from GenBank (absent in RefSeq) and found a few with wGRR higher than 0.15 for four of them (pLP39, pBS32, phiCmus, pSAM1) (suppl. table 6)). No similar phages were found for members of PiSa, cp32 and the two small communities, pp_phaeo and pp_Blicheniformis that were isolated from bacterial species with no or only a very few known phages (*Borellia, Piscirickettsia* and *Phaeobacter species*, and *Bacillus licheniformis*). In four communities P-Ps were only identified from the phage database (two large and two small communities), typically from bacterial clades where partition and replication functions are poorly known. Only 59 of the 780 P-Ps were outside communities, most being singletons (n=47) with very low wGRR to other P-Ps (Fig. S3). One prominent singleton is the crAssphage, where we could identify significant matches to HMM profiles specific for plasmid-like replication genes (previously reported in (67)). So far, no lysogenic module or integrase genes were reported for Crass-like *Bacteroides* phages, but co-replication with the host was previously described for at least one member of the crass-like phages (68), fitting the definition of P-P.

**Figure 3:**
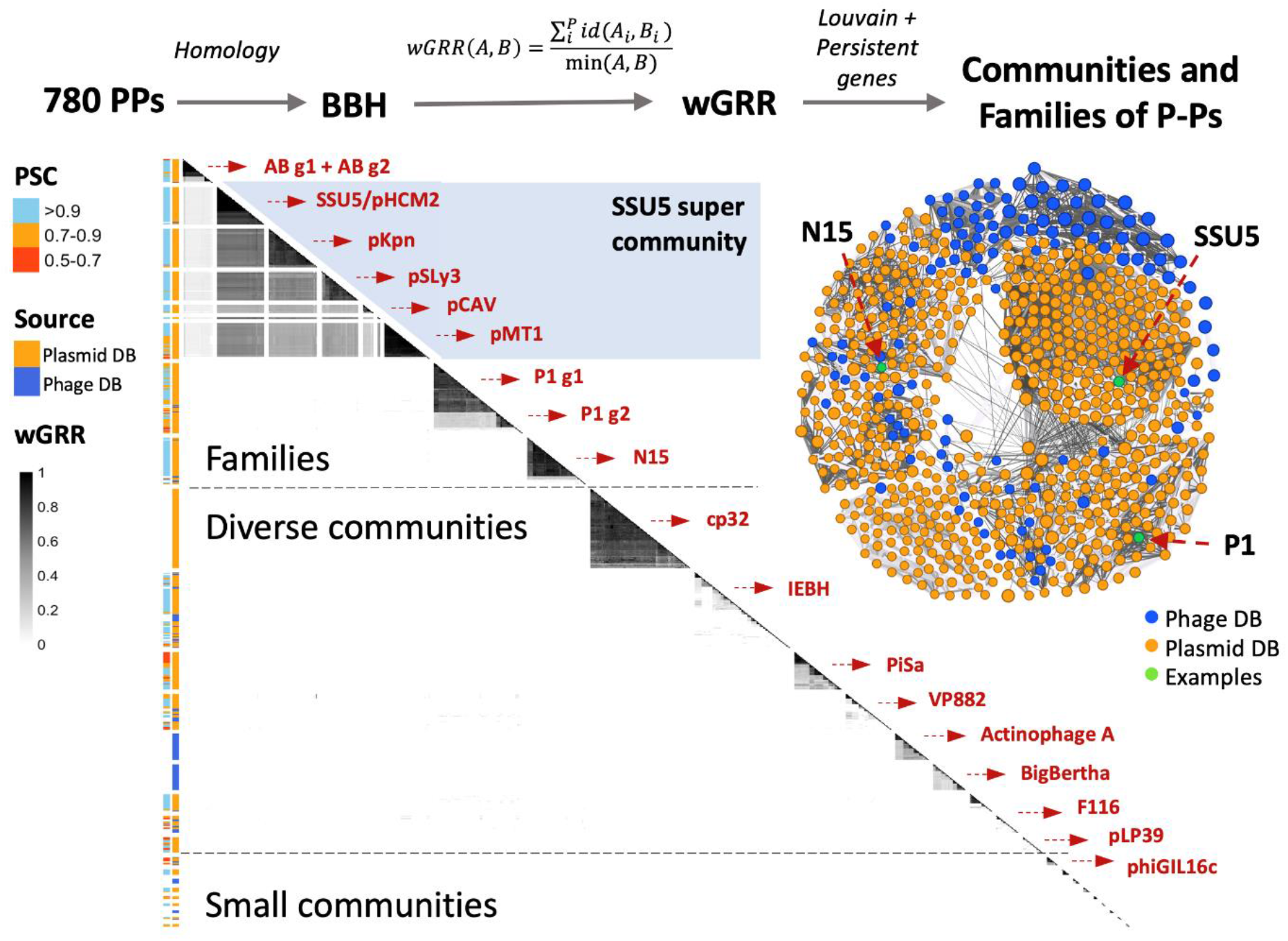
Sequence similarity network and detected communities. The communities are separated by gaps for better visibility. They were extracted, ordered in the figure by hierarchical clustering, and named after a representative P-P or a bacterial clade (in red). In the one-sided heatmap (below main diagonal), each row represents a P-P (n = 721). The 59 P-Ps not in communities were excluded (see Fig. S3). The range of the wGRR is given by the grey scale bar (from white to black). The first column on the left of the heatmap shows the phage score (PSC, given by the random forest models) and the second column indicates the database where the P-P was identified. The graph of the wGRR matrix is displayed on the right side of the heatmap. Communities that were curated into families are shown above.

We used the wGRR values and the pangenomes of the communities to curate the large communities with high average wGRR values into homogenous groups, that we call families (see Methods, Fig. 3, Fig. S4AB). The curation process yielded eight P-P families of which two, the P1 and AB, were further split into subfamilies (P1-g1, P1-g2 and AB-g1, AB-g2). Most of the members of a family are hosted by closely related bacteria, e.g. those of the AB family are from *Acinetobacter*, of the pMT1 family are from *Yersinia pestis* and those of the pKpn family are from *Klebsiella* (suppl. table 8). Overall, 42% of the P-Ps can be classed in a small number of families. The remaining elements are in communities of very diverse P-Ps and will require further data to be curated.

### Are P-Ps more like phages or more like plasmids?

It is usual to class phages according to their virion structure and plasmids in groups of replication incompatibility (Inc types). P-Ps can be classed relatively to phage taxonomy and plasmid incompatibility groups, because they encode virions and plasmid replicases. We used the virus taxonomic information from the NCBI to class the P-Ps identified in the phage database. Those identified in the plasmid database lack such information and we predicted their virus taxonomy using random forest models (Fig. S5C, see Methods and Text S1 for details). The vast majority (97%) of the assigned P-Ps are *Siphoviridae* (SSU5 superfamily and N15 family) and *Myoviridae* (P1, Fig. 4A). We could not predict a reliable virus taxonomic classification for 25.9% of the P-Ps identified in the plasmid database (suppl. table 4). Overall, the assignment of a defined P-P family to a virus taxonomical group was consistent, i.e. P-Ps from the same family usually had similar classifications. But in some highly diverse communities, those that we could not curate into families, there are sometimes different virus taxonomic groups (suppl. table 8). This confirms the need to acquire further information on these communities before proceeding to their classification into families.

**Figure 4:**
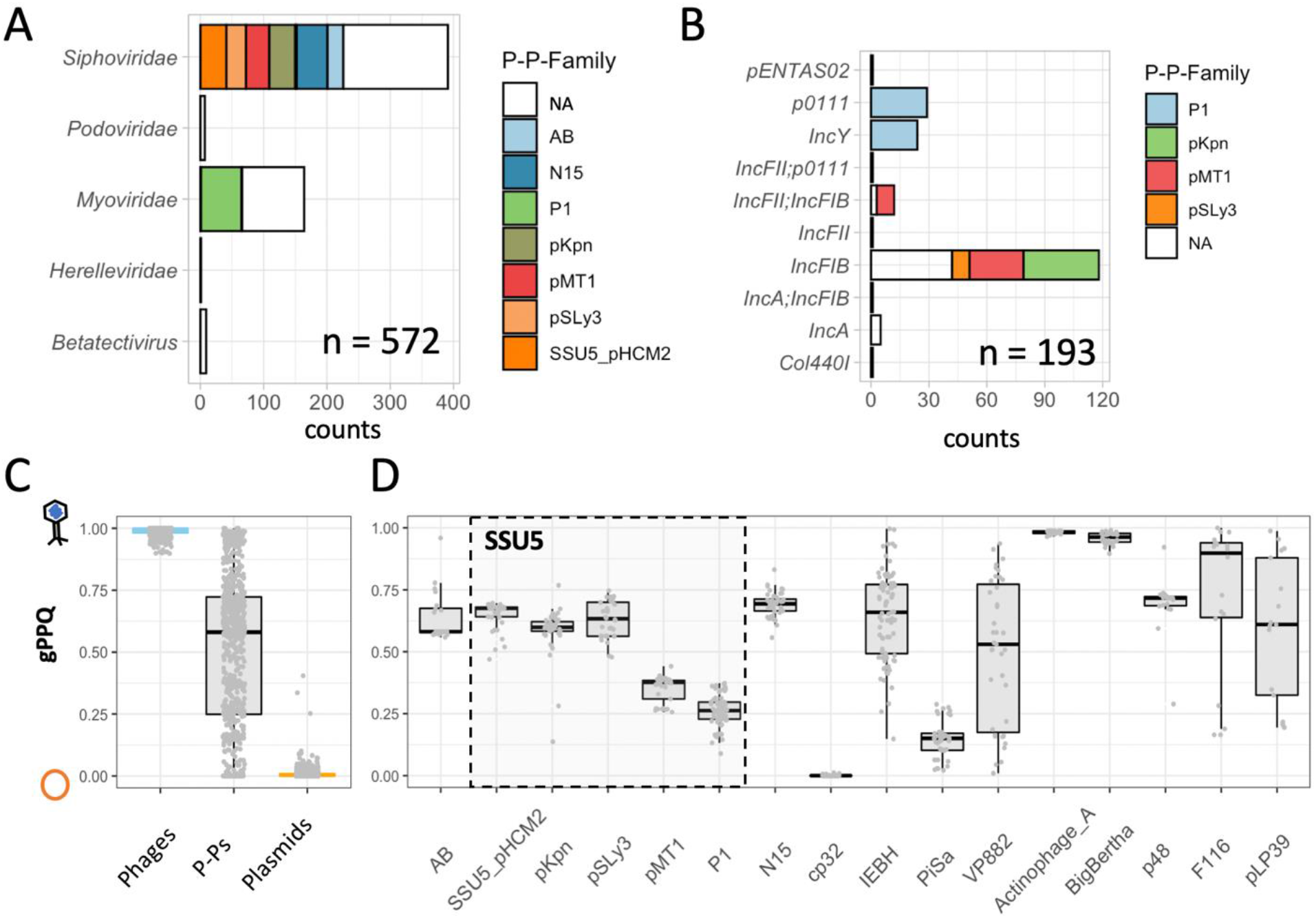
Classification of P-Ps relative to phages and plasmids. **A** and **B**. Distribution of P-Ps in terms of virus taxonomy and of incompatibility types. NA: non-curated communities. **C**. Boxplots of the genomic phage-plasmid quotients (gPPQs) for P-Ps (n=677, grey) (suppl. table 4), phages (n=458, blue) and plasmids (n=1121, orange). A few P-Ps contained only a few genes homologous to phage or plasmid genomes. To increase the accuracy of the analysis, only elements with more than 10 genes with a PPQ were considered (see Methods). **D**. Same as C for the P-P families (AB to N15) or communities (the rest) with at least 10 elements.

The Inc types of P-Ps were predicted using PlasmidFinder (57). Note that most P-Ps could not be typed. This is somewhat expected, since the PlasmidFinder database is much more detailed for *Enterobacteriaceae* than for other clades (57) and even in the remaining well-studied Proteobacteria most plasmids cannot be typed (69). When families could be systematically typed, they tended to reveal only one or two types. Notably, most P-Ps (130/193) were from the IncFIB group mainly represented by members of the SSU5 super community (Fig. 4B).

P-Ps are both phages and plasmids. Yet, from a functional and evolutionary point of view, it is interesting to address the question whether they are more like phages or more like plasmids. To answer this question, we quantified how many of their genes are homologous to those of plasmids or phages using a score that we termed the phage-plasmid quotient (PPQ). This is the number of homologs to phages divided by the number of homologs to phages and plasmids (see Methods). Its average across the P-P, termed gPPQ, ranges from 0 (only plasmid homologs) to 1 (only phage homologs). We compared the gPPQ scores of P-Ps (suppl. table 4) with those of a control group consisting of 458 phages and 1121 plasmids with similar size and host distribution as the P-Ps (Fig. S6A). Expectedly, the values for plasmids are systematically close to zero whereas those of phages are always close to one. In contrast, P-Ps have intermediate values dispersed between 0 and 1 (Fig. 4C). When these values are analyzed within each family, their dispersion decreases, showing that within-family variation is smaller than between-family variation. These values also tend to be slightly higher than 0.5, indicating the presence of more homologs to phages than to plasmids (Fig 4D). The analysis of non-curated communities shows a more diverse picture, where some P-Ps are systematically more like phages, such as the BigBertha and Actinophage A clusters and others tend to be more like plasmids (e.g. PiSa or the cp32) (Fig. 4D). The latter finding is consistent with the inability of our model to predict phage functions in the cp32 elements. The non-curated communities with most heterogeneous values of gPPQ (suppl. table 4) tend to correspond to those with low mean_wGRR_ values within the community (Fig 3, suppl. table 7). This is consistent with existence of heterogeneous types of P-Ps in these communities and explains why they could not be curated into homogeneous families.

Overall, these results show that P-Ps have many traits of phages and plasmids, and most curated families have slightly more phage-associated than plasmid-associated genes. This is not wholly unexpected, since the number of genes minimally required for a dsDNA phage is much higher than the one required for a non-conjugative plasmid. These results raise the question of the type and level of conservation of non-essential phage and plasmid genes in the families or communities of P-Ps. To answer this question, we computed the pangenome of each group of P-Ps and identified the genes present in most elements (persistent genes), present in very few (cloud genes) and the others (shell genes, see Methods). These analyses are addressed in the next sections.

### The P1 family has two distinct subfamilies

The P1 community was curated by removing seven P-Ps with few persistent genes and low wGRRs with the other members. It was then split using the wGRR and the family pangenome data into two subfamilies (Fig. S9A), where the larger one includes P1 (P1-g1) and the smaller subfamily (P1-g2) contains members that are closely related to D6, a known P1-related but GenBank-based P-P (9). The separation between the two subfamilies is clear, since the average within-subfamilies wGRRs is ∼0.75 and the one between subfamilies is only 0.23 (Fig 5B, Fig. S9C). In addition, the distinction between some of the elements of the two groups was previously described (9). The persistent genomes of the subfamilies are comparable in size (P1-g1:77 vs P1-g2:61) and split into six conserved regions separated by clusters of shell and cloud genes (Fig. S9B). In spite of the conservation of the genetic organization between the subfamilies, the one including P1 has 2.1x more shell genes and 3.6x more cloud genes than the other subfamily, suggesting that it is more plastic (or more ancient).

**Figure 5:**
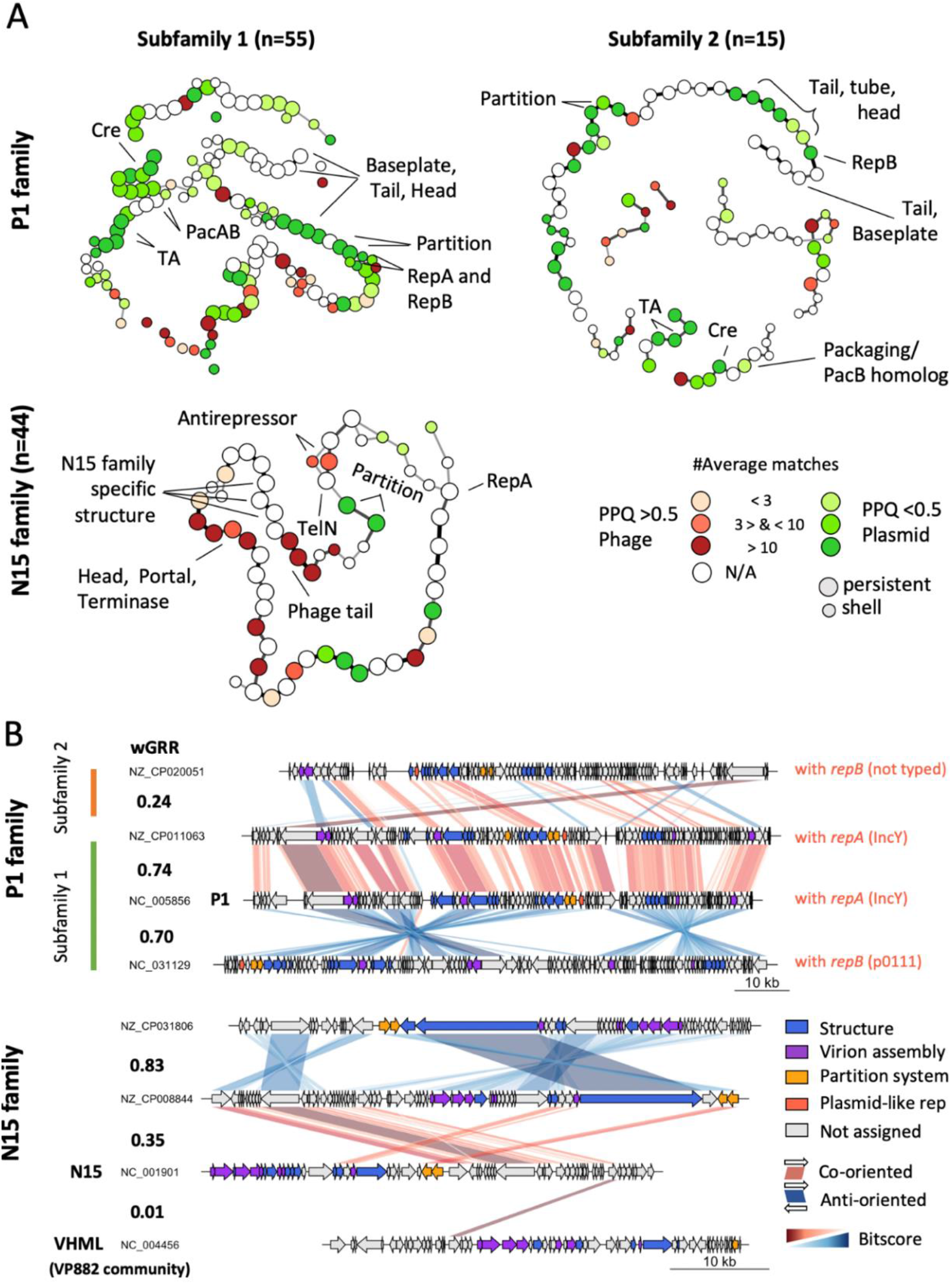
Comparative genome analysis of the P1 and N15 family. A. Pangenome graphs of the N15 and P1 families (the latter is split into two sub-families). The nodes represent genes of the persistent or shell genomes (see Figure S8C and Figure S9B for the entire pangenomes). The node colors indicate the phage-plasmid quotient (PPQ) scores (red for phage- and green for plasmid-association) that are computed from the average number of matches of the gene family with phage and plasmid genomes. The edges indicate contiguity between two genes in the P-P and their thickness indicates the frequency of this contiguity. For clarity, we removed the edges when the neighborhood was very rare: for N15<25%, for P1<15%. B. Comparisons between selected replicons plotted using genoplotR (70). Similarity between co-oriented bi-directional best hits (BBH) is shown in red and between anti-oriented ones in blue. Color intensity reflects the degree of gene similarity. The values of wGRR are shown between the pairs of elements.

The gPPQ of the P1 community tends to be smaller than 0.5 (Fig 4D), because there are more persistent gene families associated to plasmid sequences than to phages (Fig. 5A). Among these, one finds the typical plasmid core functions, but also a well-known toxin-antitoxin system (*doc, phd*) and the Cre recombinase. In terms of genetic organization, the partition and replication systems are co-localized in subfamily 1 and separated by 14 persistent genes in the other. Interestingly, subfamily 1 elements encode either a *repA* or a *repB* replicase, but never both. These genes are very different, but they are found at the same position (Fig. 5A), between two co-linear blocks that are in inverted orientation in each subfamily (Fig. 5B). These differences fit the Inc type classification, since P-Ps with *repA* are IncY and those with *repB* are p0111 (Fig. 4B, Fig. 5A). This suggests that P-Ps from both families can be maintained in a single host as plasmids, because they are compatible in terms of replication.

Even if genes homologous to phages tend to be less abundant than those homologous to plasmids, we found many persistent genes involved in the phage lytic cycle (holins, terminases, tails, baseplate proteins). Counterintuitively, some genes that are usually associated to phage functions (tails, phage head, tube proteins), have more often homologs in plasmids than in phages, resulting in a gene family with a PPQ lower than 0.5 (Fig. 5A). We assume that the causative plasmids are either defective (unrecognizable) P-Ps or plasmids that acquired structural phage genes by recombination. We also found that the *pacAB genes*, encoding the two subunits of the P1 terminase, are conserved only in the P1-g1 subfamily. The second subfamily only has *pacB* (Fig. 5A). This suggests that general transducers like P1 are more likely to be found within the P1-group. In contrast, the *phd*/*doc* TA system is highly conserved between the two subfamilies (Fig. S9C).

### N15 family is widely spread in *Enterobacteria* and characterized by the presence of the telomerase

The N15 family (n=44) was built from the N15 community by removing P-Ps (n=7) with small wGRRs between them and with the other elements ((Fig. 5A, Fig S8A-B). Most P-Ps are found in *Klebsiella* genomes (n=41), two in *E. coli* and one in *Citrobacter freundii* (suppl. table 2). Linear *Vibrio* phages from the VP882 community (such as VP882, VHML and phiHAP-1) were reported to genome organizations similar to N15 (24). This is confirmed by our analysis (Fig. 5B). However, the wGRR values between these two communities are very low (wGRR<0.01, Fig S8D) and this led us to class them apart. The genome size of the N15 family is comprised between 46.4 kb and 82.0 kb (median_N15_ = 55.3 kb). Its pangenome graph reveals the existence of three syntenic arrays separated by three small variable clusters of shell genes (Fig S8C). The *telN* gene family encoding the protelomerase is needed to maintain a linear genome. It is present in all P-Ps of the family. This strongly suggests that all these elements have linear replicons. One should note that many of the GenBank files identify these replicons as circular, but we did not find published evidence of this. Given the ubiquity of the protelomerase in the family, these are probably erroneous annotations. The partition systems, the telomerase and the *repA* gene families are present and co-localized in most of the genomes, confirming that they are defining traits of the family.

The phage-associated functions in the N15 P-Ps are more numerous than those of plasmids and also tend to be co-localized in the replicon. Some of these genes seem specific of the family (minor tails, tail tubes) whereas others have homologs in other phages (encoding head, capsid and tail proteins) (Fig. 5A, suppl. table 8). The majority of the latter are from phages infecting Enterobacteria, e.g. phage HK225 and phi80, but there are some homologs among *Burkholderia* or *Pseudomonas* phages (suppl. table 9). Two gene families, one in the persistent and the other in the shell genome, encode alternative SOS-dependent phage anti-repressors homologous to those of some lambdoid phages. They are located in the same genomic region, but they are never present in the same genome, and are very similar (79% identity covering ∼99% of the sequence) suggesting that they are fast-evolving orthologs (Fig. 5A).

### The AB family is specific to Acinetobacter

The curation of this community led to the exclusion of two distantly related members, resulting in the AB-family that contains only P-Ps from *Acinetobacter spp*. It is noteworthy, that one of the excluded members is the phage RhEph10 of Rhizobium that is homologous to the known P-P pLM21S1 of *Sinorhizobium* Rhizobium (71). The AB family is the only family/superfamily lacking (to the best of our knowledge) a known phage. A screening of the GenBank phage database revealed two phages, the *Klebsiella* phage ST13-OXA48phi12.3 and the *Pesudomonas* phage Nickie, that are distantly related to the family (highest wGRR: 0.18 and 0.15) (supp. table 5). The family was further split into two subfamilies AB-g1 (n=19) and AB-g2 (n=5) with similar replicon sizes (∼110 kb) (Fig S10A), but showing low overall similarity (wGRR = 0.25, Fig. 6A, Fig S10C). We found 54 persistent genes conserved across the two subfamilies showing that even if the percent similarity of proteins is low, both subfamilies share a large number of homologs (Fig. 6B, Fig S10C). They include many phage-related functions such as terminases, tail, assembly, head, but not lysozymes. It is noteworthy that some of these are homologous to tail proteins of phages from *Enterobacteria* and *Burkholderia* (suppl. table 12 and 13). Key functions involved in homologous recombination (*recA* and *recF*), and in partition are homologous in the two subfamilies. The latter includes ParB encoding genes (involved in DNA segregation) that occur in two copies in most of the elements (Fig. 6A). In spite of these commonalities, the pangenome of subfamily 1 is larger than the one of subfamily 2, especially in what concerns the shell and cloud genomes (Fig. S10B). Also, the plasmid-related functions - replication and *parA* partition gene - are highly divergent (Fig. 6AB, Fig S10C). In summary, the AB subfamilies are relatively small and found mostly in *A. baumannii* strains (only 2 P-P are found in other species). Like for the N15 and P1 families, the genes homologous to phages tend to be in the persistent genome, whereas the plasmid-associated genes are more diverse and variable. As described below, the AB family also shares similarities with the SSU5 super-community.

**Figure 6:**
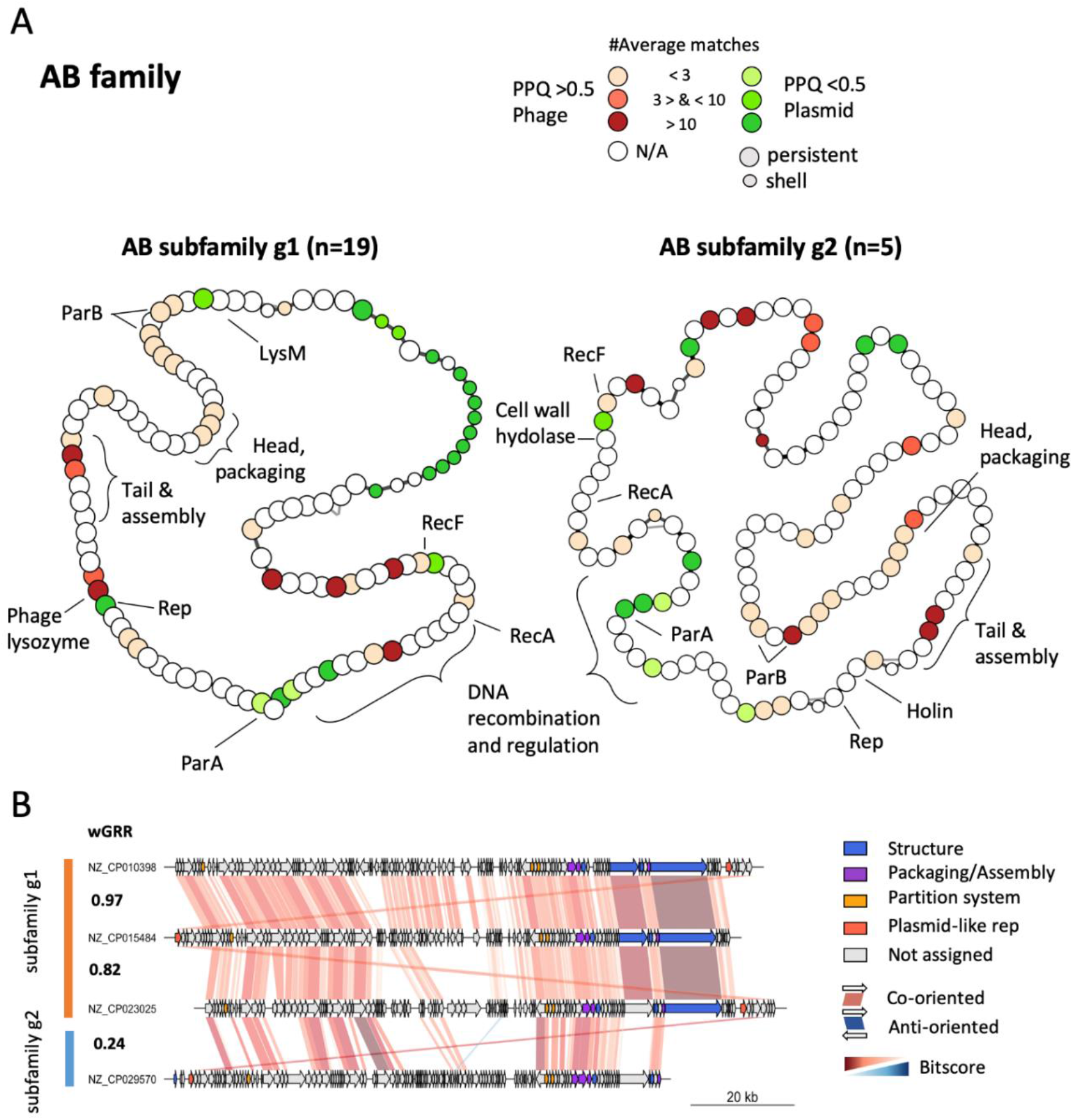
Pangenome analysis of the AB family. A. Pangenome graphs of the two subfamilies of the AB family. B. Comparisons between selected replicons. For details, see legend of Figure 5.

### The SSU5 supercommunity is the largest group of P-Ps

The SSU5 supercommunity includes the five communities SSU5_pHCM2 (n=41), pKpn (n=42), pSLy3 (n=32), pMT1 (n=39), pCAV (n=9) and two other P-Ps (Fig. 3). All these elements are related to SSU5 and to each other (average wGRR between communities in the range 0.23 to 0.59) (Fig. S10C). They were isolated from different enterobacterial hosts, including *E. coli, K. pneumonia, S. enterica* and *Y. pestis*. The curation process of the supercommunity led to the exclusion of three elements among the pMT1 and pSLy3 communities and the entire pCAV community resulting in the SSU5 superfamily. The pCAV family was excluded because it has only a few persistent genes in common with the rest (Fig. S11A). Hence, the SSU5 superfamily consists of four families of P-Ps (SSU5_pHCM2, pKpn, pSLy3, pMT1) (Fig. S11, S12). The SSU5 superfamily has a complex and large pangenome consisting of 35 persistent, 281 shell and 815 cloud gene families (Fig. S11B). In addition, in the shell genome (genes present at intermediate frequencies in the superfamily) some genes are present in multiple, but not all, P-P families (Fig. S11A). This suggests the existence of genetic flux across these families of P-Ps.

We detected more phage-like genes (n=16) than plasmid-like ones (n=3) in the persistent genome of the superfamily. Interestingly, most of the former are clustered in one region, denominated the phage-array, with many homologs in the pCAV family (Fig. 7A). These genes encode phage tails, capsids and terminases. As found in the N15 and AB families, the tail genes are related to genes found in lambdoid *Siphoviridae* from *Enterobacteria* (suppl. table 18). Most of the persistent gene families that are not in the phage-array are involved in DNA recombination e.g. resolvases, tyrosine-recombinases and RecA-like proteins (Fig. 7A). In contrast, the plasmid-like genes are much more abundant in the shell and especially in the cloud genomes where they are more than 4.3 times more frequent than phage homologs (Fig. S7A, suppl. table. 18). Although the function of most gene families of the shell genome is not known, they also include Toxin-Antitoxin and restriction modification systems, anti-restriction mechanism (such as ArdA-like proteins) and putative virulence factors like pili assembling proteins (PapC and PapD) (72). The cloud genome of the supercommunity is very large (>2500 gene families), reflecting the diversity of these P-Ps (suppl. table 18).

**Figure 7:**
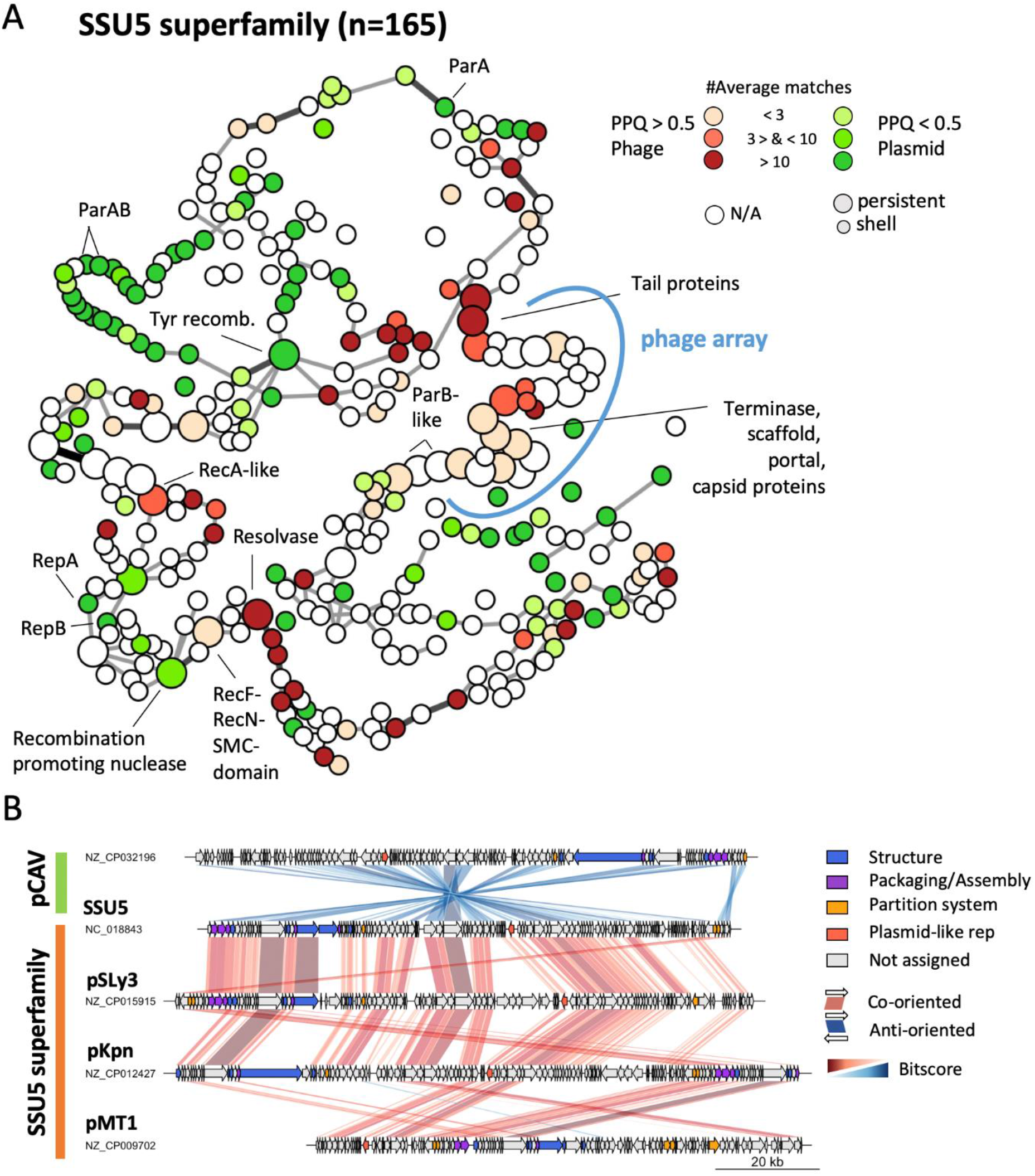
Conserved patterns in genomes of the SSU5 superfamily. A. Pangenome graph of the SSU5 superfamily. B. Comparisons between selected replicons. For details, see legend of Figure 5. The family pCAV was excluded from the analysis of the pangenome because it’s not included in the superfamily (see main text).

The comparison of the superfamily’s pangenome with those of each family revealed conserved regions beyond the abovementioned arrays of genes for phage structural proteins and recombination functions (Fig. 8). Some of these regions are specific to a particular family (blue nodes in Fig. 8) whereas others are conserved across different families or even at the level of the whole superfamily (orange/yellow nodes in Fig. 8). Most notably, many of the P-Ps from the pMT1 family share a specific set of co-localized plasmid-like gene (Fig 8, blue nodes in pMT1). These genes are not found in the other three P-P families (pSLy3, pKpn and SSU5_pHCM2) which pangenomes, however, show a more similar organization, more intersections in the graphs and more common frequent genes (Fig. 8). Nevertheless, this does not translate into larger differences in terms of genetic plasticity, since the pangenome and wGRR matrices of the pKpn and pSLy3 families show higher diversity of gene repertoires than those of pMT1 and SSU5_pHCM2 families (Fig S12BC, Fig S13BC). Interestingly, the SSU5 superfamily shows also some similarity to the AB-family (wGRR_mean_ = 0.08) (Fig. S14A). Most of these homologous genes are found in the persistent genome of the SSU5 superfamily, especially in the arrays of genes encoding the phage structural genes and the recombinases (Fig. S14B). Hence, the SSU5 superfamily, the pCAV and the AB family are evolutionarily related, especially concerning phage and recombination functions. As most families, the SSU5 superfamily has a core of conserved and co-localized phage-like genes that accounts for a large fraction of the persistent genes of pangenomes and a larger number of plasmid-like genes that differ more widely both within and between the families.

**Figure 8:**
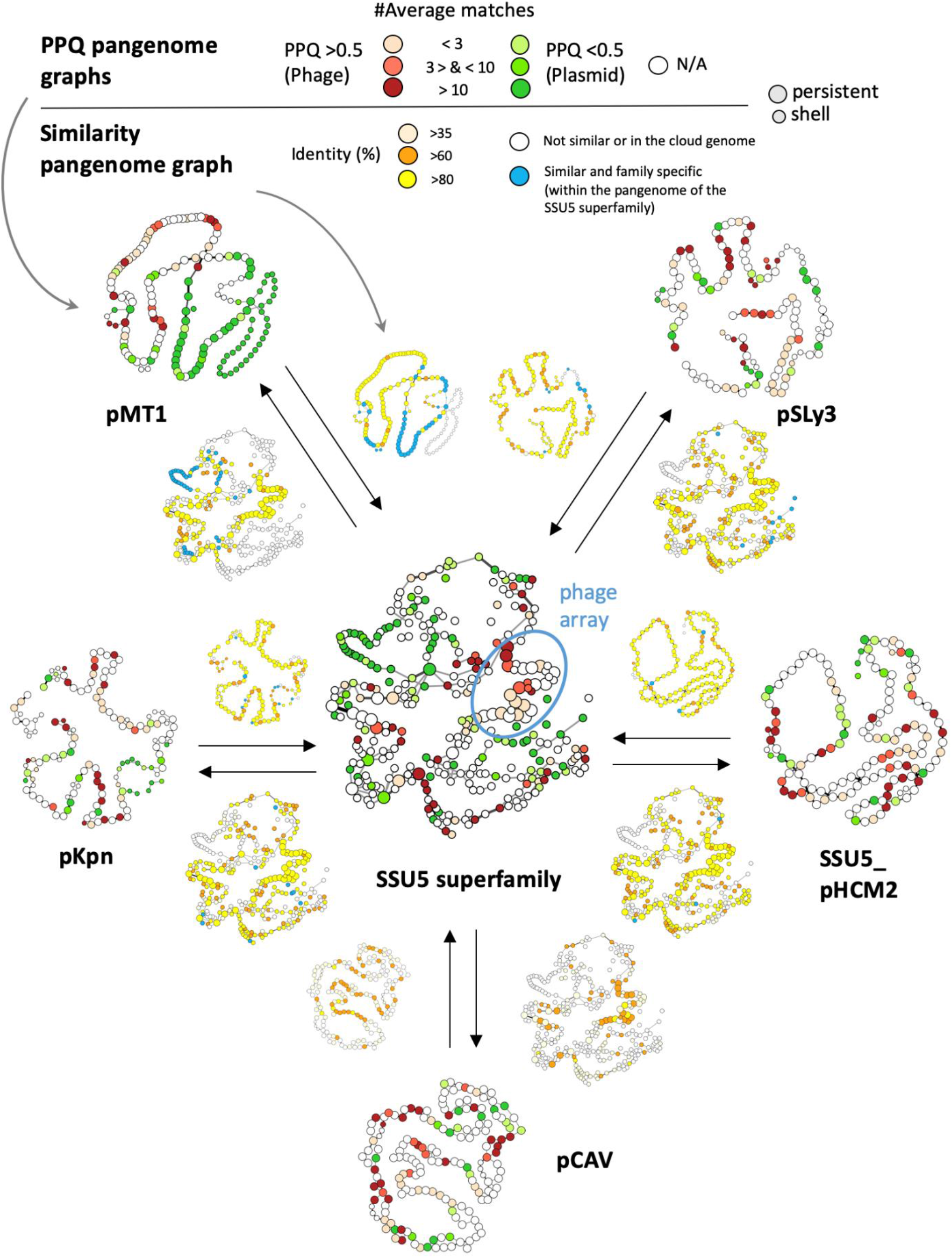
Analysis of the SSU5 superfamily and the pCAV family. Pangenome graphs of the SSU5_pHCM2, pKpn, pMT1, pSLy3 and pCAV families and the SSU5 superfamily were colored in function of the values of PPQ (larger graphs) and similarity to the pangenome of the SSU5 superfamily (smaller graphs next to the arrows). Nodes and edges are as in Fig 5. The average number of homologs of a gene family with phage and/ or plasmid genomes is given in the PPQ graphs. Genes that are specific to one family are shown in blue in the SSU5 similarity graphs. Otherwise, genes and their orthologues (BBH) found in at least two families are indicated in orange/yellow/light yellow nodes (depending on their average identity) (see Methods). An example: The pMT1 pangenome (top left) is highly related to the SSU5 superfamily (center), since the two similarity pangenome graphs next to the arrows show many similarities (colored in light yellow, orange to yellow). However, some co-localized gene families are only found in the pMT1 family (they are indicated in blue).

### Non-curated P-P communities

#### IEBH

This large community includes the known P-Ps IEBH (73), and was mostly isolated from *Bacilli* and *Clostridia*. The replicons are very diverse (mean wGRR = 0.06) and their level of similarity is extremely variable (coefficient of variation (cv) =267%) (suppl. table 7). Their average genome size is 49 kb, but the range of sizes is very large (from 16 kb to 160 kb) (suppl. table 4).

#### phiGIL16c

This small community from *Bacillus* includes phiGIL16c, which was shown to form phage particles and be maintained as a linear plasmid (27). All community members are assigned to the Betatectiviruses, showing that P-Ps can be found in dsDNA phages outside Caudovirales. Five of the nine replicons are annotated in RefSeq as linear including phiGIL16c, Bam35c and pBClin15 (27, 62). The genomes range between 13 kb and 15 kb in size (suppl. table 4) and are closely related (mean wGRR = 0.59) (suppl. table 7). Our screening failed to identify partition or replication systems in these P-P, suggesting that plasmid maintenance uses so far unknown mechanisms. As a result, all these elements were identified in the phage database.

#### VP882

The P-Ps of this community are too diverse to be put in large families (mean wGRR = 0.1) (suppl. table 7) and their sizes are extremely variable (from 16.6 kb to 241.8 kb (suppl. table 4), average = 40.9 kb). They are found across Proteobacteria, including *Vibrio* (VP882) (26), *Arsenophonus, Cupriavidus, Halomonas, Burkholderia* (KS-14) (74), *Klebsiella* or *Escherichia* (P88) (supplemental table 4). It is noteworthy that P88 was isolated from a lysogenic *E. coli* strain after induction (75). Our screening identifies partition and plasmid-like replication genes, but P88 was previously found to be integrated (75) suggesting that it may have episomal and integrative states. Several of the *Vibrio* and *Halomonas* P-Ps have been reported to have linear replicons (26). However, the protelomerase is present in only five P-Ps, suggesting that most of the elements have circular replicons.

#### BigBertha

This heterogeneous community of P-Ps from *Bacillus* has large replicons (average size 159.7 kb) (suppl. table 8). They were all identified in the phage database, and many were previously described as strictly virulent and belonging to the SPO1-like phages (76–78) (suppl. table 4). However, all of them had homologs of the partition systems of the IEBH group, which contains a *bona fide* P-P. Since no clear plasmid state was reported and it was suggested that the partition genes might be involved in host sporulation (79), we are not very confident that this community is constituted of P-Ps.

#### cp32

These plasmids from *Borellia*/*Borreliella* are around 30 kb in size and are quite similar (mean wGRR = 0.72, cv = 17%) (Fig. S4). They were previously proposed to be P-Ps (64), and one related member (phiBB-1) was experimentally proven to form virions (65). Although phiBB-1 was not sequenced, its genome hybridizes with cp32 DNA (80) and it was demonstrated that it can transduce cp32s (65). However, cp32 elements score poorly in our random forest models (PSC between 0.003 and 0.375) and we found very few proteins with phage homologs (PPQ between 0 and 0.012) (suppl. table. 4). Moreover, our search in the GenBank database revealed no phage homolog (with at least wGRR >= 0.1). Since the plasmids of *Borrelia* have been described as recombining very frequently (81), many cp32 may be defective P-Ps.

#### pLP39

In this diverse community with 17 members, 14 of them were isolated from *Lactobacilli*. Members of the community are poorly related (wGRR < 0.11, suppl. table 7). Their sizes vary between 19.7 kb and 108.3 kb with an average of 40.0 kb (suppl. table 4 and 8). So far, none of them were experimentally reported to be P-Ps. However, our models predict a high PSC >0.9 for nine P-Ps (suppl. table 4) and we could find homologous phages, such as phage Sha1 and PM411 (wGRR 0.39 and 0.76) (suppl. table 6), suggesting the pLP39 community contains true P-Ps.

#### Actinophage_A

These P-Ps were identified from the actinophages of the cluster A. They were known to encode partition systems, lack integration cassettes and remain extrachromosomal (25, 82). Some elements infect *Gordonia terrae*, but the majority infects *Mycobacterium smegmatis*. Their sizes are quite similar (average of 52.9 kb) (suppl. table 8), even if their gene repertoires are only moderately related (mean wGRR = 0.42) (suppl. table 7).

#### PiSa

This heterogeneous community was identified exclusively from plasmids of *Piscirickettsia salmonis*. Their sizes vary widely from 31.9 kb to 188.3 kb and their gene repertoires are moderately related (mean wGRR = 0.42). There is no experimental evidence that any of its members forms phage particles and, we could not find relatives in the GenBank phage database. In addition, since the phage scores for 37% of the members were relatively low (PSC < 0.7) (suppl. table 4), it is possible that some of these elements have lost part of the phage genes. This is consistent with previous observations that *Piscirickettsia* plasmids are highly mosaic due to a suspected high activity of transposases (83).

#### F116

This highly diverse community includes the known F116 P-P of *Pseudomonas aeruginosa* (63). The replicons are poorly related (average wGRR = 0.11) (suppl. table 7), their sizes range widely from 21.6 to 243.8 kb (suppl. table 4), and their taxonomic classification is inconsistent within the community (suppl. table 8). We could not detect a partition or plasmid replication system in F116, whereas most other elements of the community encoded at least a ParA (including Phages SE1, ST160 and phi297). In two of the other members of the community, D3 and phiSG1 (suspected but not proven to be P-Ps (84, 85)), we found homologs to plasmid replicases. The *Pseudomonas* phage YMC11/02/R656 also encodes a plasmid replicase. Eleven P-Ps were identified among plasmids of *Klebsiella* and *Shigella*. Although no experiments proved them to be P-P, some show very high phage score (PSC>0.9) (suppl. table 4).

## Conclusion

The 780 putative P-Ps identified in this work represent 5.6% of the phage and 7.3% of the plasmid database. This shows that P-Ps are a significant fraction of elements classed as phages or plasmids. But these precise numbers should be taken with care, since they vary widely across clades and depend on several factors. First, we assume that we can identify phage and plasmid associated functions. This is probably true for most key phage functions in the best studied bacterial clades, but may not be true for phage and plasmid functions in Spirochaetes, Bacteroides and other clades. This will result in an underestimate of the number of P-Ps, especially when searching for plasmid functions in phages, because replicases evolve fast and many known plasmids lack recognizable replicases (6, 57). Second, we cannot ascertain if these P-Ps are functional. Bacterial chromosomes contain many defective prophages (86, 87), and this may also be the case of some P-Ps. Similarly, elements identified from the phage database may have lost plasmid functions (although loss of both replicase and partition systems will make them undetectable in our screen). Third, some elements may oscillate between integrated and extra-chromosomal states (8), blurring the distinction between chromosomal prophages and P-Ps. Fourth, some putative P-Ps may be prophages integrated in plasmids. We expect prophage integration in plasmids to be rare, since phages are thought to select for highly conserved integration sites in chromosomes (plasmids tend to be present in few strains of a species). To minimize this problem, we excluded megaplasmids and secondary chromosomes from the analyses. Finally, one cannot exclude the possibility that some putative P-Ps are actually caught in a relation of pseudolysogeny with the bacterial host (88) (although that would leave unexplained the presence of plasmid-like functions). In spite of these caveats, the observation that the best-studied clades in the database (enterobacteria and *Bacillus*) have more P-Ps than the average Bacteria and that these have clear similarities to known P-Ps, suggests that we have underestimated the number of P-Ps.

By clustering P-Ps in terms of similarity of gene repertoires, we showed that P-Ps can be classed in distinct groups (communities or homogenous families). While the persistent genes between different communities were usually very divergent, we could systematically identify homologs in key phage functions across them. For example, the AB, pCAV and SSU5 families have homologous persistent genes, suggesting a distant evolutionary association between these families. Hence, P-Ps are probably ancient, i.e. they are not just transient chimeric mobile elements recently created from recombination between phages and plasmids. Further work on the very heterogeneous communities of P-Ps that remained non-curated may reveal yet novel families of these elements and will facilitate future studies on the evolutionary associations between families.

Intriguingly, most tailed P-Ps in our dataset are *Siphoviridae* or *Myoviridae*, and few are *Podoviridae*. The reasons for this are unclear, but the database does over-represent the first two classes of tailed phages (89). P-Ps can also be found in *Tectiviridae* opening the possibility of their presence in other types of poorly characterized phages. *Inoviridae* are known to replicate actively without inducing the lytic cycle, in which they resemble plasmids and P-Ps (90, 91). However, these ssDNA phages replicate while actively producing and exporting virions, explaining why we chose to exclude them from this analysis. Hybrids between viruses and plasmids have also been reported in archaea (92, 93), suggesting that some archaeal viruses are also P-Ps. Metagenomics based studies are uncovering many novel phage genomes and it will be interesting to assess how many of these are P-Ps.

P-Ps are phages and plasmids. Hence, one expects them to carry accessory traits from both. Indeed, some P-P families have many homologs to phage genes, whereas others tend to have more homologs in plasmids. The study of the pangenomes revealed that phage homologs tend to be more conserved than plasmid homologs. In contrast, the latter tend to be more frequent in variable regions. As a result, even if there are on average more phage than plasmid homologs in P-Ps, the latter are more variable and may thus account for a large fraction of the genes providing adaptive phenotypes to bacterial hosts.

## DATA AVAILABILITY

All genomes were taken from public databases. The necessary data are provided in the article and in the supplemental material. Any further requests e.g. on data processing can be sent to.

## FUNDING

This work was funded by a grant from the ANR Labex IBEID 10-LABX-0062 for EP and from SALMOPROPHAGE ANR-16-CE16-0029 for JMS. The laboratory is supported by grants from the INCEPTION project (PIA/ANR-16-CONV-0005) and the Fédération pour la Recherche Médicale [Equipe FRM/EQU201903007835].

## CONFLICT OF INTEREST

The authors declare that there is no conflict of interest.

## ACKNOWLEDGEMENTS

The authors would like to thank Olaya Rendueles-Garcia, Antoine Frenoy and Charles Coluzzi for comments and suggestions. Moreover, many thanks to Jean Cury, Sophie Abby and Bertrand Néron for providing useful tools such as MacSyFinder and a pipeline for annotating plasmid functions.

## TABLE LEGENGS

Supplemental table 1

**Phage specific HMM profiles group1**. HMM profiles associated with phages from the pVOG, PFAM and TIGRFAM databases (DB) annotated and classed into functional categories. The column Phage_profile_cluster indicates the cluster composed of multiple homologous HMM profiles identified by profile-profile alignment.

Supplemental table 2

**Phage specific HMM profiles group2**: pVOG profiles with a VQ >0.75 and based on more than 15 protein sequences. The table indicates the number of proteins identified (and the number of different genomes), the annotation and functional categories and the viral quotient data.

Supplemental table 3

**Plasmid specific HMM profiles**

Partition systems protein profiles from PFAM, TIGFRAM, pVOG and eggNOG databases to detect plasmid-related functions in phages (the others were retrieved from (94)).

Supplemental table 4

**List of phage-plasmids identified in this study**. P-Ps are listed with their general features: source of the P-P (plasmid or phage database), NCBI accession name and id, number of genes and genome size, host genus, PSC mean and standard deviation, assignment to (super) community/ (super-/, sub-) family, (predicted) virus taxonomy, incompatibility type, gPPQ score and number of considered genes. The last column indicates references to works showing that the elements are P-Ps or have traits typical of P-Ps (like plasmid partition genes in phages).

Supplemental table 5

**wGRR similarity table of phage-plasmids** (based on the single NCBI accession IDs).

Supplemental table 6

List of phages from the virus NCBI database that have a homology (wGRR > 0.1) to P-Ps which are in communities where all elements were identified in the plasmid database (no element from the Virus database in RefSeq). NCBI accessions of the P-P and the phage, as well as information on the size, the wGRR, the number of BBHs, the P-P community and the host of the phage are indicated.

Supplemental table 7

**wGRR similarity table of P-P groups**. Group sizes of P-P groups, mean, standard deviation and coefficient of variation of the wGRR similarity between different P-P groups.

Supplemental table 8

**Summary of the characteristics of P-P (super)communities and (sub)families:** group type, group name, group size, average genome size, average number of genes per genome, average PSC and PPQ scores, assigned virus and host taxonomy (numbers in brackets indicate the counts).

Supplemental table 9 to 19 (organized in the same way)

**Overview on the gene families of the P-P (sub-/ super-) families**.

Category of the gene family (persistent, shell and cloud). The names of the gene families are based on the NCBI accession IDs of the first genes. The NCBI IDs of all genes that were used for a gene family are indicated in the NCBI_Member column. Database used to annotate the genes (PFAM, TIGFRAM, pVOG and eggNOG (Viruses and bactNOG profiles only)). The IDs represent the indexes that are found in the indexed pangenome graphs (Figure S15 to S19). Characterization of the gene families in terms of numbers, origin, number of hits to plasmids and phages and average PPQ. The numbers in square brackets (next to the name) show how many gene families have BBHs to genes of the affected phage/ plasmid. Since the PPQ may differ from gene to gene within a gene family, the average PPQ was used. Moreover, the average number of hits to phages and plasmids were used and the NCBI names and NCBI accessions of the first 500 phages and plasmids are listed.

Explanatory example: Gene family phiKO2p16 (ID=76), a gene family within the N15 pangenome, is annotated to encode minor tail genes and is based on 42 genes. On average 31 phages and plasmids have BBHs with genes from phiKO2p16. The average PPQ is 0.99 and is considered as phage-like. The highest number of BBHs to plasmids is two that have the NCBI accession NC_013856 and NC_015062. The number 41 (in squared brackets) behind NC_015062, show that this plasmid (from *Rahnella sp*. Y9602) has overall 41 BBHs with the pangenome of the N15 family. In addition, the highest number of BBHs to phage genomes is 39.

**Supplemental table 9: Gene families of the N15 family**.

**Supplemental table 10: Gene families of the P1 subfamily1**.

**Supplemental table 11: Gene families of the P1 subfamily2**.

**Supplemental table 12: Gene families of the AB subfamily1**.

**Supplemental table 13: Gene families of the AB subfamily2**.

**Supplemental table 14: Gene families of the SSU5_pHCM2 family**.

**Supplemental table 15: Gene families of the pKpn family**.

**Supplemental table 16: Gene families of the pSLy3 family**.

**Supplemental table 17: Gene families of the pMT1 family**.

**Supplemental table 18: Gene families of the SSU5 superfamily**.

**Supplemental table 19: Gene families of the pCAV family**.

## Notes

### Competing Interest Statement

The authors have declared no competing interest.

## REFERENCES

1. Frost, L.S., Leplae, R., Summers, A.O. and Toussaint, A. (2005) Mobile genetic elements: the agents of open source evolution. Nat. Rev. Microbiol., 3, 722–732.

2. Touchon, M., Moura de Sousa, J.A. and Rocha, E.P. (2017) Embracing the enemy: the diversification of microbial gene repertoires by phage-mediated horizontal gene transfer. Curr. Opin. Microbiol., 38, 66–73.

3. Chiang, Y.N., Penadés, J.R. and Chen, J. (2019) Genetic transduction by phages and chromosomal islands: The new and noncanonical. PLoS Pathog, 15, e1007878.

4. Smillie, C., Garcillán-Barcia, M.P., Francia, M.V., Rocha, E.P.C. and de la Cruz, F. (2010) Mobility of Plasmids. Microbiol Mol Biol Rev, 74, 434–452.

5. Gandon, S. (2016) Why Be Temperate: Lessons from Bacteriophage λ. Trends Microbiol., 24, 356–365.

6. Cury, J., Oliveira, P.H., de la Cruz, F. and Rocha, E.P.C. (2018) Host Range and Genetic Plasticity Explain the Coexistence of Integrative and Extrachromosomal Mobile Genetic Elements. Mol Biol Evol, 35, 2230–2239.

7. Łobocka, M.B., Rose, D.J., Plunkett, G., Rusin, M., Samojedny, A., Lehnherr, H., Yarmolinsky, M.B. and Blattner, F.R. (2004) Genome of Bacteriophage P1. J. Bacteriol, 186, 7032–7068.

8. Utter, B., Deutsch, D.R., Schuch, R., Winer, B.Y., Verratti, K., Bishop-Lilly, K., Sozhamannan, S. and Fischetti, V.A. (2014) Beyond the Chromosome: The Prevalence of Unique Extra-Chromosomal Bacteriophages with Integrated Virulence Genes in Pathogenic Staphylococcus aureus. PLOS ONE, 9, e100502.

9. Gilcrease, E.B. and Casjens, S.R. (2018) The genome sequence of Escherichia coli tailed phage D6 and the diversity of Enterobacteriales circular plasmid prophages. Virology, 515, 203–214.

10. Ravin, N.V., Svarchevsky, A.N. and Dehò, G. (1999) The anti-immunity system of phage-plasmid N15: identification of the antirepressor gene and its control by a small processed RNA. Mol. Microbiol., 34, 980–994.

11. Tabassum Khan, N. (2017) Mechanisms of Plasmid Replication. J Proteomics Bioinform, 10, 211–213.

12. Salje, J. (2010) Plasmid segregation: how to survive as an extra piece of DNA. Crit. Rev. Biochem. Mol, 45, 296–317.

13. Sengupta, M. and Austin, S. (2011) Prevalence and Significance of Plasmid Maintenance Functions in the Virulence Plasmids of Pathogenic Bacteria. Infect Immun, 79, 2502–2509.

14. Ravin, N.V. (2011) N15: The linear phage–plasmid. Plasmid, 65, 102–109.

15. Lindler, L.E., Plano, G.V., Burland, V., Mayhew, G.F. and Blattner, F.R. (1998) Complete DNA Sequence and Detailed Analysis of the Yersinia pestis KIM5 Plasmid Encoding Murine Toxin and Capsular Antigen. Infect Immun, 66, 5731–5742.

16. Venturini, C., Zingali, T., Wyrsch, E.R., Bowring, B., Iredell, J., Partridge, S.R. and Djordjevic, S.P. (2019) Diversity of P1 phage-like elements in multidrug resistant Escherichia coli. Sci. Rep, 9, 18861.

17. Bertani, G. (1951) STUDIES ON LYSOGENESIS I.: The Mode of Phage Liberation by Lysogenic Escherichia coli. J. Bacteriol, 62, 293–300.

18. Lennox, E.S. (1955) Transduction of linked genetic characters of the host by bacteriophage P1. Virology, 1, 190–206.

19. Skorupski, K., Sauer, B. and Sternberg, N. (1994) Faithful Cleavage of the P1 Packaging Site (pac) Requires Two Phage Proteins, PacA and PacB, and Two Escherichia coli Proteins, IHF and HU. J. Mol. Biol., 243, 268–282.

20. Yarmolinsky, M. and Hoess, R. (2015) The Legacy of Nat Sternberg: The Genesis of Cre-lox Technology. Annu. Rev. Viro, 2:1, 25–40.

21. McLellan, M.A., Rosenthal, N.A. and Pinto, A.R. (2017) Cre-loxP-Mediated Recombination: General Principles and Experimental Considerations. Curr. Protoc. mouse Biol, 7, 1–12.

22. Ravin, N.V. (2015) Replication and Maintenance of Linear Phage-Plasmid N15. Microbiol. Spectr., 3, (1):PLAS-0032-2014.

23. Knott, S.E., Milsom, S.A. and Rothwell, P.J. (2019) The Unusual Linear Plasmid Generating Systems of Prokaryotes. In Bacteriophages - Perspectives and Future. IntechOpen.

24. Hammerl, J.A., Jäckel, C., Funk, E., Pinnau, S., Mache, C. and Hertwig, S. (2016) The diverse genetic switch of enterobacterial and marine telomere phages. Bacteriophage, 6, e1148805.

25. Dedrick, R.M., Mavrich, T.N., Ng, W.L., Cervantes Reyes, J.C., Olm, M.R., Rush, R.E., Jacobs-Sera, D., Russell, D.A. and Hatfull, G.F. (2016) Function, expression, specificity, diversity and incompatibility of actinobacteriophage parABS systems. Mol. Microbiol., 101, 625–644.

26. Lan, S.-F., Huang, C.-H., Chang, C.-H., Liao, W.-C., Lin, I.-H., Jian, W.-N., Wu, Y.-G., Chen, S.-Y. and Wong, H. (2009) Characterization of a New Plasmid-Like Prophage in a Pandemic Vibrio parahaemolyticus O3:K6 Strain. Appl. Environ. Microbiol., 75, 2659–2667.

27. Verheust, C., Fornelos, N. and Mahillon, J. (2005) GIL16, a new gram-positive tectiviral phage related to the Bacillus thuringiensis GIL01 and the Bacillus cereus pBClin15 elements. J. Bacteriol., 187, 1966–1973.

28. Myers, G.S.A., Rasko, D.A., Cheung, J.K., Ravel, J., Seshadri, R., DeBoy, R.T., Ren, Q., Varga, J., Awad, M.M., Brinkac, L.M., et al. (2006) Skewed genomic variability in strains of the toxigenic bacterial pathogen Clostridium perfringens. Genome Res, 16, 1031–1040.

29. Kim, M., Kim, S. and Ryu, S. (2012) Complete Genome Sequence of Bacteriophage SSU5 Specific for Salmonella enterica serovar Typhimurium Rough Strains. J. Virol., 86, 10894–10894.

30. Kim, M., Kim, S., Park, B. and Ryu, S. (2014) Core Lipopolysaccharide-Specific Phage SSU5 as an Auxiliary Component of a Phage Cocktail for Salmonella Biocontrol. Appl Environ Microbiol, 80, 1026–1034.

31. Octavia, S., Sara, J. and Lan, R. (2015) Characterization of a large novel phage-like plasmid in Salmonella enterica serovar Typhimurium. FEMS Microbiol Lett, 362, 1–9.

32. Kidgell, C., Pickard, D., Wain, J., James, K., Diem Nga, L.T., Diep, T.S., Levine, M.M., O’Gaora, P., Prentice, M.B., Parkhill, J., et al. (2002) Characterisation and distribution of a cryptic Salmonella typhi plasmid pHCM2. Plasmid, 47, 159–171.

33. Falgenhauer, L., Yao, Y., Fritzenwanker, M., Schmiedel, J., Imirzalioglu, C. and Chakraborty, T. (2014) Complete Genome Sequence of Phage-Like Plasmid pECOH89, Encoding CTX-M-15. Genome Announc, 2, e00356–14.

34. Yang, L., Li, W., Jiang, G.-Z., Zhang, W.-H., Ding, H.-Z., Liu, Y.-H., Zeng, Z.-L. and Jiang, H.-X. (2017) Characterization of a P1-like bacteriophage carrying CTX-M-27 in Salmonella spp. resistant to third generation cephalosporins isolated from pork in China. Sci Rep, 7, 40710: 1–8.

35. Santamaría, R.I., Bustos, P., Sepúlveda-Robles, O., Lozano, L., Rodríguez, C., Fernández, J.L., Juárez, S., Kameyama, L., Guarneros, G., Dávila, G., et al. (2014) Narrow-Host-Range Bacteriophages That Infect Rhizobium etli Associate with Distinct Genomic Types. Appl Environ Microbiol, 80, 446–454.

36. Hammerl, J.A., Klein, I., Appel, B. and Hertwig, S. (2007) Interplay between the Temperate Phages PY54 and N15, Linear Plasmid Prophages with Covalently Closed Ends. J Bacteriol, 189, 8366–8370.

37. O’Leary, N.A., Wright, M.W., Brister, J.R., Ciufo, S., Haddad, D., McVeigh, R., Rajput, B., Robbertse, B., Smith-White, B., Ako-Adjei, D., et al. (2016) Reference sequence (RefSeq) database at NCBI: current status, taxonomic expansion, and functional annotation. Nucleic Acids Res., 44, D733–745.

38. Hatcher, E.L., Zhdanov, S.A., Bao, Y., Blinkova, O., Nawrocki, E.P., Ostapchuck, Y., Schäffer, A.A. and Brister, J.R. (2017) Virus Variation Resource - improved response to emergent viral outbreaks. Nucleic Acids Res, 45, D482–D490.

39. R Core Team (2020) R: A language and environment for statistical computing. R Foundation for Statistical Computing, Vienna, Austria.

40. Eddy, S.R. (2011) Accelerated Profile HMM Searches. PLoS Comput. Biol., 7, e1002195.

41. El-Gebali, S., Mistry, J., Bateman, A., Eddy, S.R., Luciani, A., Potter, S.C., Qureshi, M., Richardson, L.J., Salazar, G.A., Smart, A., et al. (2019) The Pfam protein families database in 2019. Nucleic Acids Res, 47, D427–D432.

42. Haft, D.H., Loftus, B.J., Richardson, D.L., Yang, F., Eisen, J.A., Paulsen, I.T. and White, O. (2001) TIGRFAMs: a protein family resource for the functional identification of proteins. Nucleic Acids Res, 29, 41–43.

43. Huerta-Cepas, J., Szklarczyk, D., Heller, D., Hernández-Plaza, A., Forslund, S.K., Cook, H., Mende, D.R., Letunic, I., Rattei, T., Jensen, L.J., et al. (2019) eggNOG 5.0: a hierarchical, functionally and phylogenetically annotated orthology resource based on 5090 organisms and 2502 viruses. Nucleic Acids Res., 47, D309–D314.

44. Grazziotin, A.L., Koonin, E.V. and Kristensen, D.M. (2017) Prokaryotic Virus Orthologous Groups (pVOGs): a resource for comparative genomics and protein family annotation. Nucleic Acids Res, 45, D491–D498.

45. Abby, S.S., Néron, B., Ménager, H., Touchon, M. and Rocha, E.P.C. (2014) MacSyFinder: A Program to Mine Genomes for Molecular Systems with an Application to CRISPR-Cas Systems. PLOS ONE, 9, e110726.

46. Fouts, D.E. (2006) Phage_Finder: Automated identification and classification of prophage regions in complete bacterial genome sequences. Nucleic Acids Res, 34, 5839–5851.

47. Söding, J. (2005) Protein homology detection by HMM–HMM comparison. Bioinformatics, 21, 951–960.

48. Blondel, V.D., Guillaume, J.-L., Lambiotte, R. and Lefebvre, E. (2008) Fast unfolding of communities in large networks. J. Stat. Mech., 2008, P10008.

49. Cury, J., Touchon, M. and Rocha, E.P.C. (2017) Integrative and conjugative elements and their hosts: composition, distribution and organization. Nucleic Acids Res., 45, 8943–8956.

50. Cury, J., Abby, S.S., Doppelt-Azeroual, O., Néron, B. and Rocha, E.P.C. (2020) Identifying Conjugative Plasmids and Integrative Conjugative Elements with CONJscan. Methods Mol. Biol., 2075, 265–283.

51. Arndt, D., Grant, J.R., Marcu, A., Sajed, T., Pon, A., Liang, Y. and Wishart, D.S. (2016) PHASTER: a better, faster version of the PHAST phage search tool. Nucleic Acids Res, 44, W16–W21.

52. Wright, M.N. and Ziegler, A. (2017) ranger: A Fast Implementation of Random Forests for High Dimensional Data in C++ and R. J. Stat. Softw, 77, 1–17.

53. Robin, X., Turck, N., Hainard, A., Tiberti, N., Lisacek, F., Sanchez, J.-C. and Müller, M. (2011) pROC: an open-source package for R and S+ to analyze and compare ROC curves. BMC Bioinformatics, 12, 77.

54. Steinegger, M. and Söding, J. (2017) MMseqs2 enables sensitive protein sequence searching for the analysis of massive data sets. Nat. Biotechnol., 35, 1026–1028.

55. Bobay, L.-M., Rocha, E.P.C. and Touchon, M. (2013) The Adaptation of Temperate Bacteriophages to Their Host Genomes. Mol Biol Evol, 30, 737–751.

56. Christensen, A.P. (2018) NetworkToolbox: Methods and Measures for Brain, Cognitive, and Psychometric Network Analysis in R. R J., 10, 422–439.

57. Carattoli, A., Zankari, E., García-Fernández, A., Voldby Larsen, M., Lund, O., Villa, L., Møller Aarestrup, F. and Hasman, H. (2014) In silico detection and typing of plasmids using PlasmidFinder and plasmid multilocus sequence typing. Antimicrob. Agents Chemother., 58, 3895–3903.

58. Gautreau, G., Bazin, A., Gachet, M., Planel, R., Burlot, L., Dubois, M., Perrin, A., Médigue, C., Calteau, A., Cruveiller, S., et al. (2020) PPanGGOLiN: Depicting microbial diversity via a partitioned pangenome graph. PLOS Computational Biology, 16, e1007732.

59. Bastian, M., Heymann, S. and Jacomy, M. (2009) Gephi: an open source software for exploring and manipulating networks. In Third international AAAI conference on weblogs and social media.

60. Csardi, G. and Nepusz, T. The igraph software package for complex network research. InterJournal, Complex Systems, 1695.

61. Zhang, C., Feng, Y., Liu, F., Jiang, H., Qu, Z., Lei, M., Wang, J., Zhang, B., Hu, Y., Ding, J., et al. (2017) A Phage-Like IncY Plasmid Carrying the mcr-1 Gene in Escherichia coli from a Pig Farm in China. Antimicrob Agents Chemother, 61, e02035–16.

62. Strömsten, N.J., Benson, S.D., Burnett, R.M., Bamford, D.H. and Bamford, J.K.H. (2003) The Bacillus thuringiensis linear double-stranded DNA phage Bam35, which is highly similar to the Bacillus cereus linear plasmid pBClin15, has a prophage state. J. Bacteriol., 185, 6985–6989.

63. Miller, R.V., Pemberton, J.M. and Clark, A.J. (1977) Prophage F116: Evidence for Extrachromosomal Location in Pseudomonas aeruginosa Strain PAO. J Virol, 22, 844–847.

64. Casjens, S.R., Gilcrease, E.B., Vujadinovic, M., Mongodin, E.F., Luft, B.J., Schutzer, S.E., Fraser, C.M. and Qiu, W.-G. (2017) Plasmid diversity and phylogenetic consistency in the Lyme disease agent Borrelia burgdorferi. BMC Genomics, 18, 165.

65. Eggers, C.H., Kimmel, B.J., Bono, J.L., Elias, A.F., Rosa, P. and Samuels, D.S. (2001) Transduction by ϕBB-1, a Bacteriophage of Borrelia burgdorferi. J Bacteriol, 183, 4771–4778.

66. Wetzel, K.S., Aull, H.G., Zack, K.M., Garlena, R.A. and Hatfull, G.F. (2020) Protein-Mediated and RNA-Based Origins of Replication of Extrachromosomal Mycobacterial Prophages. mBio, 11, e00385–20.

67. Dutilh, B.E., Cassman, N., McNair, K., Sanchez, S.E., Silva, G.G.Z., Boling, L., Barr, J.J., Speth, D.R., Seguritan, V., Aziz, R.K., et al. (2014) A highly abundant bacteriophage discovered in the unknown sequences of human faecal metagenomes. Nat. Commun., 5, 1–11.

68. Shkoporov, A.N., Khokhlova, E.V., Fitzgerald, C.B., Stockdale, S.R., Draper, L.A., Ross, R.P. and Hill, C. (2018) ΦCrAss001 represents the most abundant bacteriophage family in the human gut and infects Bacteroides intestinalis. Nat. Commun., 9, 4781.

69. Shintani, M., Sanchez, Z.K. and Kimbara, K. (2015) Genomics of microbial plasmids: classification and identification based on replication and transfer systems and host taxonomy. Front Microbiol, 6, 242:1–16.

70. Guy, L., Roat Kultima, J. and Andersson, S.G.E. (2010) genoPlotR: comparative gene and genome visualization in R. Bioinformatics, 26, 2334–2335.

71. Dziewit, L., Pyzik, A., Szuplewska, M., Matlakowska, R., Mielnicki, S., Wibberg, D., Schlüter, A., Pühler, A. and Bartosik, D. (2015) Diversity and role of plasmids in adaptation of bacteria inhabiting the Lubin copper mine in Poland, an environment rich in heavy metals. Front Microbiol, 6, 152:1–12.

72. Allen, W.J., Phan, G. and Waksman, G. (2012) Pilus biogenesis at the outer membrane of Gram-negative bacterial pathogens. Curr. Opin. Struct. Biol., 22, 500–506.

73. Smeesters, P.R., Drèze, P.-A., Bousbata, S., Parikka, K.J., Timmery, S., Hu, X., Perez-Morga, D., Deghorain, M., Toussaint, A., Mahillon, J., et al. (2011) Characterization of a novel temperate phage originating from a cereulide-producing Bacillus cereus strain. Res. Microbiol., 162, 446–459.

74. Lynch, K.H., Stothard, P. and Dennis, J.J. (2010) Genomic analysis and relatedness of P2-like phages of the Burkholderia cepacia complex. BMC Genomics, 11, 599.

75. Chen, M., Zhang, L., Xin, S., Yao, H., Lu, C. and Zhang, W. (2017) Inducible Prophage Mutant of Escherichia coli Can Lyse New Host and the Key Sites of Receptor Recognition Identification. Front. Microbiol., 8, 147:1–13.

76. Lee, J.-H., Shin, H., Son, B., Heu, S. and Ryu, S. (2013) Characterization and complete genome sequence of a virulent bacteriophage B4 infecting food-borne pathogenic Bacillus cereus. Arch. Virol., 158, 2101–2108.

77. Gillis, A. and Mahillon, J. (2014) Phages Preying on Bacillus anthracis, Bacillus cereus, and Bacillus thuringiensis: Past, Present and Future. Viruses, 6, 2623–2672.

78. Klumpp, J., Lavigne, R., Loessner, M.J. and Ackermann, H.-W. (2010) The SPO1-related bacteriophages. Arch Virol, 155, 1547–1561.

79. El-Arabi, T.F., Griffiths, M.W., She, Y.-M., Villegas, A., Lingohr, E.J. and Kropinski, A.M. (2013) Genome sequence and analysis of a broad-host range lytic bacteriophage that infects the Bacillus cereus group. Virol J, 10, 48.

80. Eggers, C.H. and Samuels, D.S. (1999) Molecular evidence for a new bacteriophage of Borrelia burgdorferi. J. Bacteriol., 181, 7308–7313.

81. Casjens, S., Palmer, N., Vugt, R.V., Huang, W.M., Stevenson, B., Rosa, P., Lathigra, R., Sutton, G., Peterson, J., Dodson, R.J., et al. (2000) A bacterial genome in flux: the twelve linear and nine circular extrachromosomal DNAs in an infectious isolate of the Lyme disease spirochete Borrelia burgdorferi. Mol. Microbiol., 35, 490–516.

82. Mavrich, T.N. and Hatfull, G.F. (2019) Evolution of Superinfection Immunity in Cluster A Mycobacteriophages. mBio, 10, e00971–19.

83. Pesesky, M.W., Tilley, R. and Beck, D.A.C. (2019) Mosaic plasmids are abundant and unevenly distributed across prokaryotic taxa. Plasmid, 102, 10–18.

84. Miller, R.V., Pemberton, J.M. and Richards, K.E. (1974) F116, D3 and G101: Temperate bacteriophages of Pseudomonas aeruginosa. Virology, 59, 566–569.

85. Clark, A.J., Pontes, M., Jones, T. and Dale, C. (2007) A Possible Heterodimeric Prophage-Like Element in the Genome of the Insect Endosymbiont Sodalis glossinidius. J Bacteriol, 189, 2949–2951.

86. Asadulghani, M., Ogura, Y., Ooka, T., Itoh, T., Sawaguchi, A., Iguchi, A., Nakayama, K. and Hayashi, T. (2009) The Defective Prophage Pool of Escherichia coli O157: Prophage–Prophage Interactions Potentiate Horizontal Transfer of Virulence Determinants. PLoS Pathog, 5, e1000408.

87. Oliveira, J., Mahony, J., Hanemaaijer, L., Kouwen, T.R.H.M., Neve, H., MacSharry, J. and van Sinderen, D. (2017) Detecting Lactococcus lactis Prophages by Mitomycin C-Mediated Induction Coupled to Flow Cytometry Analysis. Front. Microbiol., 8, 1343: 1–8.

88. Cenens, W., Makumi, A., Mebrhatu, M.T., Lavigne, R. and Aertsen, A. (2013) Phage–host interactions during pseudolysogeny: Lessons from the Pid/dgo interaction. Bacteriophage, 3, e25029.

89. Ackermann, H.-W. and Prangishvili, D. (2012) Prokaryote viruses studied by electron microscopy. Arch Virol, 157, 1843–1849.

90. Krupovic, M. (2013) Networks of evolutionary interactions underlying the polyphyletic origin of ssDNA viruses. Curr Opin Virol, 3, 578–586.

91. Fauquet, C.M. (2006) The diversity of single stranded DNA viruses. Biodiversity, 7, 38–44.

92. Arnold, H.P., She, Q., Phan, H., Stedman, K., Prangishvili, D., Holz, I., Kristjansson, J.K., Garrett, R. and Zillig, W. (1999) The genetic element pSSVx of the extremely thermophilic crenarchaeon Sulfolobus is a hybrid between a plasmid and a virus. Mol. Microbiol., 34, 217–226.

93. Iranzo, J., Koonin, E.V., Prangishvili, D. and Krupovic, M. (2016) Bipartite Network Analysis of the Archaeal Virosphere: Evolutionary Connections between Viruses and Capsidless Mobile Elements. J. Virol., 90, 11043–11055.

94. Cury, J., Touchon, M. and Rocha, E.P.C. (2017) Integrative and conjugative elements and their hosts: composition, distribution and organization. Nucleic Acids Res., 45, 8943–8956.

