## supplemental material for "Bacteria have numerous phage-plasmid families with conserved phage and variable plasmid gene repertoires"

#### SUPPLEMENTAL METHODS AND ANALYSIS

**Text S1. Prediction of the virus taxonomy by machine learning. Training and evaluation of random forest models.**

Viruses are classed in taxonomical units depending on their origin (phage, plasmid) and virion morphology. The group of tailed dsDNA phages, termed *Caudovirales*, represents an order of bacterial viruses and consists of nine families. *Myoviridae*, *Siphoviridae* and *Podoviridae* are the three so far most prominent ones containing most of the phages (>60% of tailed phages belong to the *Siphoviridae*). The taxonomy of most phages was assigned by electron microscopy (1), but some remain unassigned (no virion formation reported).

127 of the 780 detected P-Ps were identified in the phage database meaning that their the virus taxonomy is known. 653 putative P-Ps are found in the plasmid database, where no experimental data on the virus taxonomy were available. To predict their virus taxonomy, we trained and used 10 random forest models. For the training, each model a dataset of 2000 randomly chosen phages (with known taxonomy, positive cases) and 2000 randomly chosen plasmids with an average phage score (PSC) < 0.1 (negative cases) (for details, see Methods). We included the negative data set to identify cases for which a prediction is not confident. The evaluation was done using 10 test data sets each consisting each of 500 phages and 1000 plasmids. Each model was evaluated by a data set that is independent from its train data set (as for the PSC prediction models). The taxonomy with the highest probability score was assigned to the P-P. Overall, 15000 predictions (10 models, each with 1500 predictions) were done of which 98.6% were positive (Fig. S5A, left panel). Since train and test datasets were defined randomly, a few replicons were classified only once (only by one model) and others multiple times (by up to 8 different models) (Fig. S5A, right panel). We calculated the mean probability score of the classification (and standard deviation) for all phages (n=822) and plasmids (n=835) that were at least three times classified showing an average of 98.8% ± 0.2% (Fig. S5B).

The models were then used to predict the virus taxonomy of the 653 P-Ps (found in the plasmid database). In 582 cases the assignment of a taxonomy was consistent with the predictions of the 10 models. In these cases a taxonomy was assigned, if the mean probability minus one standard deviation was higher than 0.5. Otherwise, no assignment was done. In the remaining 71 P-Ps multiple taxonomies were predicted and therefore the classification with the highest frequency was chosen e.g. if 9 models predicted *Myoviridae* and 1 model assigned *Siphoviridae* then *Myoviridae* was chosen. As for the consistent cases, a taxonomy was only assigned if the mean probability minus standard deviation was higher than 0.5. Otherwise, a taxonomy was not assigned.

**Text S2. Explanatory example of the PPQ and gPPQ calculation**

The PPQ is inspired by the Viral Quotient (VQ) of the pVOGs (http://dmk-brain.ecn.uiowa.edu/pVOGs/tutorial.html#) (2). It is the number of BBH of genes in P-Ps found in phages divided by the total counts (same analysis on phages and plasmids, normalised to the size of each of the two databases). The BBH is a bi-directional best hit between proteins in two different mobile elements, for which the e-value $<10^{-4}$, the sequence identity >= 35% and the alignment length covered at least 50% of each of the sequences. The use of BBHs, instead of just the number of homologs, means that each gene is only counted once, even if there are several homologs (e.g. duplications of transposable elements).

*For example*: A protein sequence has BBHs to 10 phages (out of 2375 in the database) and one to 1 a plasmid (out of 10785 in the database). The phage database is made of the RefSeq phages w/o the P-Ps (n=127). The plasmid database includes all RefSeq plasmids without the P-Ps (n=653) and w/o plasmids with an PSC between 0.1 and 0.5 (n=955) (for details see Methods). The PPQ is then:

$$PPQ = (10/2375) / ((10/2375) + (1/10785)) =$$
$$0.0042 / (0.0042+0.00001) = 0.998$$

BBH to only phages would lead to a PPQ of 1 and only to plasmids to 0.

The gPPQ is the average of the PPQ scores of all genes (restricted to comparisons with enough homologs per replicon).

*For example*: Let's consider a P-P with 20 genes. Ten genes matched only phage genomes (PPQ=1) and 5 matched only plasmid genomes (PPQ = 0). Five genes did not match enough homologs. The gPPQ is the average:

$$gPPQ = 10 / (10 + 5) = 0.67$$

If there are only matches to phage genomes, the gPPQ would be 1 and only matches to plasmids would lead to a gPPQ of 0. Note that the gPPQ was calculated only for P-Ps, plasmids and phages with at least 10 protein sequences for which one could compute a PPQ.

#### SUPPLEMENTAL FIGURES

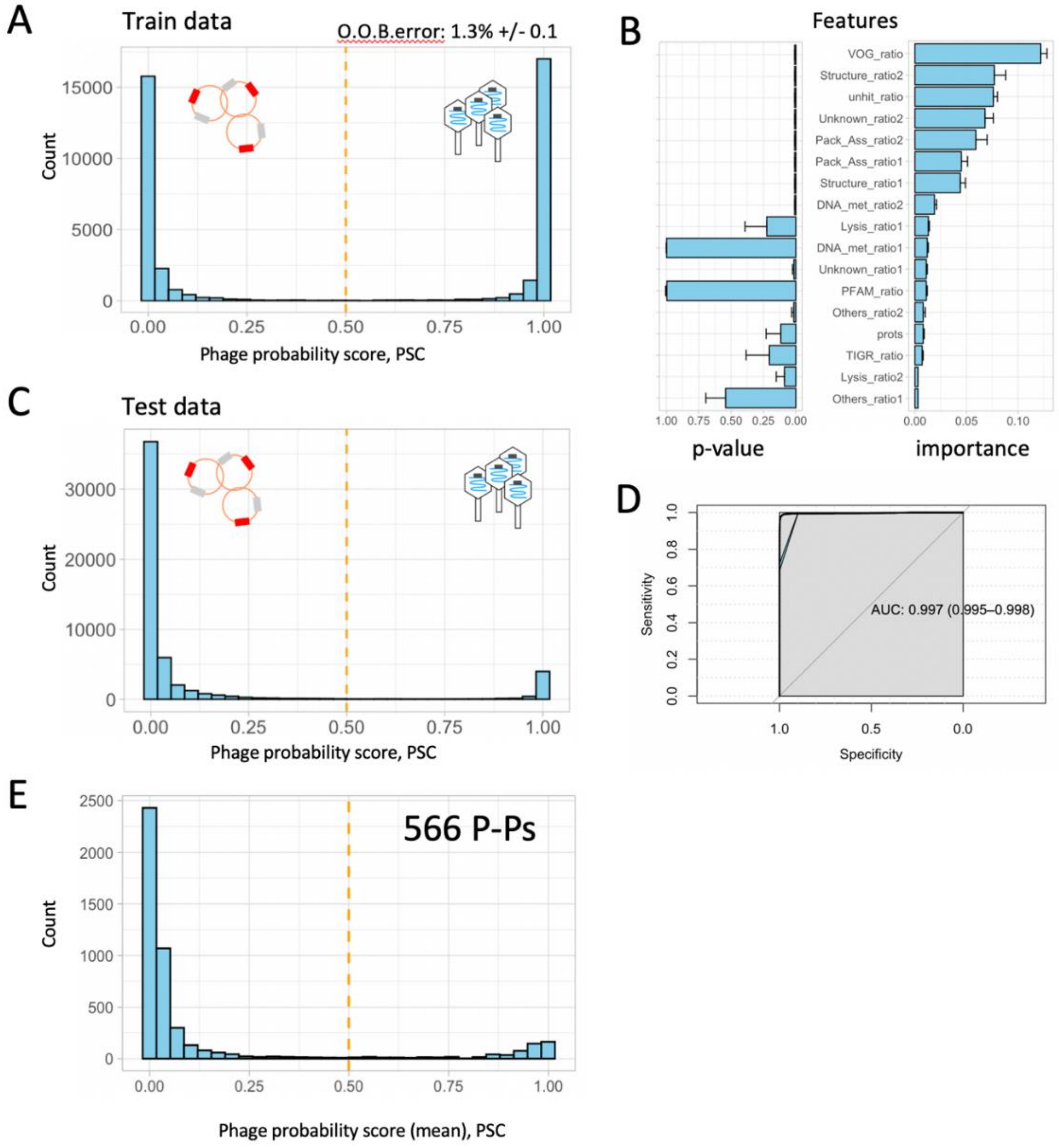

**Figure S1: Training, evaluation and application of the random forest models to predict the phage probability score (PSC).**

**A**. 10 random forest models were trained on 2000 randomly chosen phages (positives, PSC was set to 1) and 2000 randomly selected plasmids that were predicted by PHASTER (3) to not contain any phage sequences (negatives, PSC was set to 0). The mean of the out of the box (O.O.B.) error rate is 1.3% ± 0.1%. **B.** The weight of each feature (given by all models) on the decision (importance) and its p-value were calculated using the permutation option in the ranger package in R. Shown are the mean and standard deviations calculated from the 10 models for the training dataset. **C**. Evaluations of the models' classifications were done using a data set that is independent from the train data (each consisting of 4950 plasmids and 497 phages). **D.** The Area Under the Receiver Operating Characteristics were computed using the pROC package (4) in R. The confidence intervals are based on bootstraps. **E**. The 10 models were applied on plasmids that were positively predicted by PHASTER to contain prophages (including all cases: intact, questionable, incomplete). 566 plasmids with a mean phage probability larger than PSC > 0.5 were predicted (suppl. table 4).

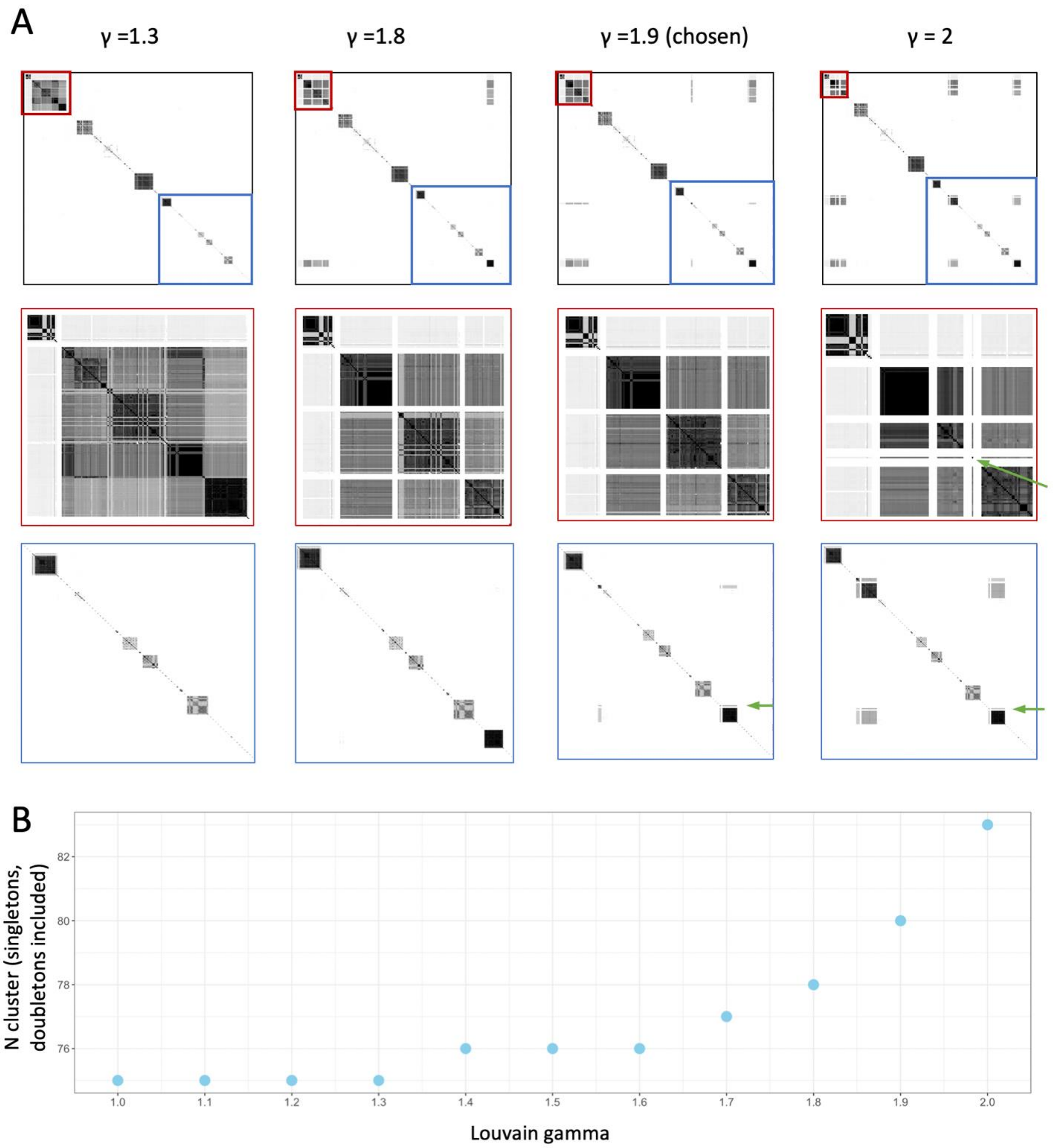

**Figure S2: Dependence of the P-P clustering on the Louvain gamma (γ) parameter.**

A. Different values for the Louvain gamma (γ) parameter were applied and evaluated using the NetworkToolbox package in R (5). Zoomed regions of the blue and red boxes are shown in the second and third row. The parameter γ = 1.9 was chosen since it resulted in a clearer clustering, especially in the largest P-P community (SSU5 super community, pointed out by the red box). In contrast, γ > 1.9 split some communities with moderate values of wGRR, resulting in too many singletons/doubletons (shown by green arrows). B. Number of communities shown for the different gamma parameters ranging from 1 to 2.

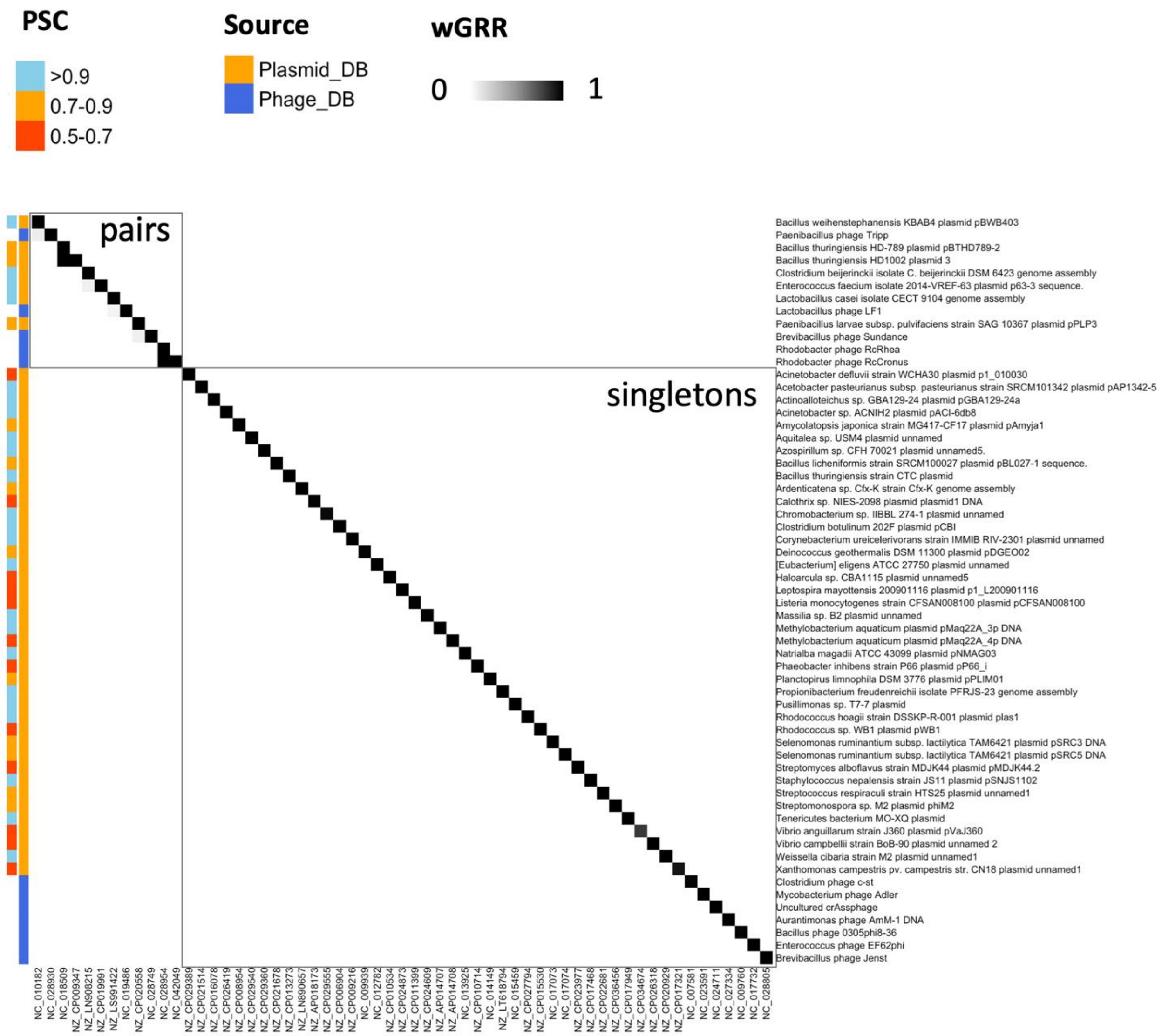

**Figure S3. wGRR matrix of 59 P-P singletons and doubletons.**

The clustering was done using the Louvain detection method (6) and resulted in 47 singletons and 12 P-Ps that are organized in pairs (doubletons). Row names are those from the NCBI database and column names are the NCBI accession numbers of the same replicon. The first column (left of the matrix) shows the mean PSC given by the prediction models and the second indicates the source of the P-P (phage or plasmid database).

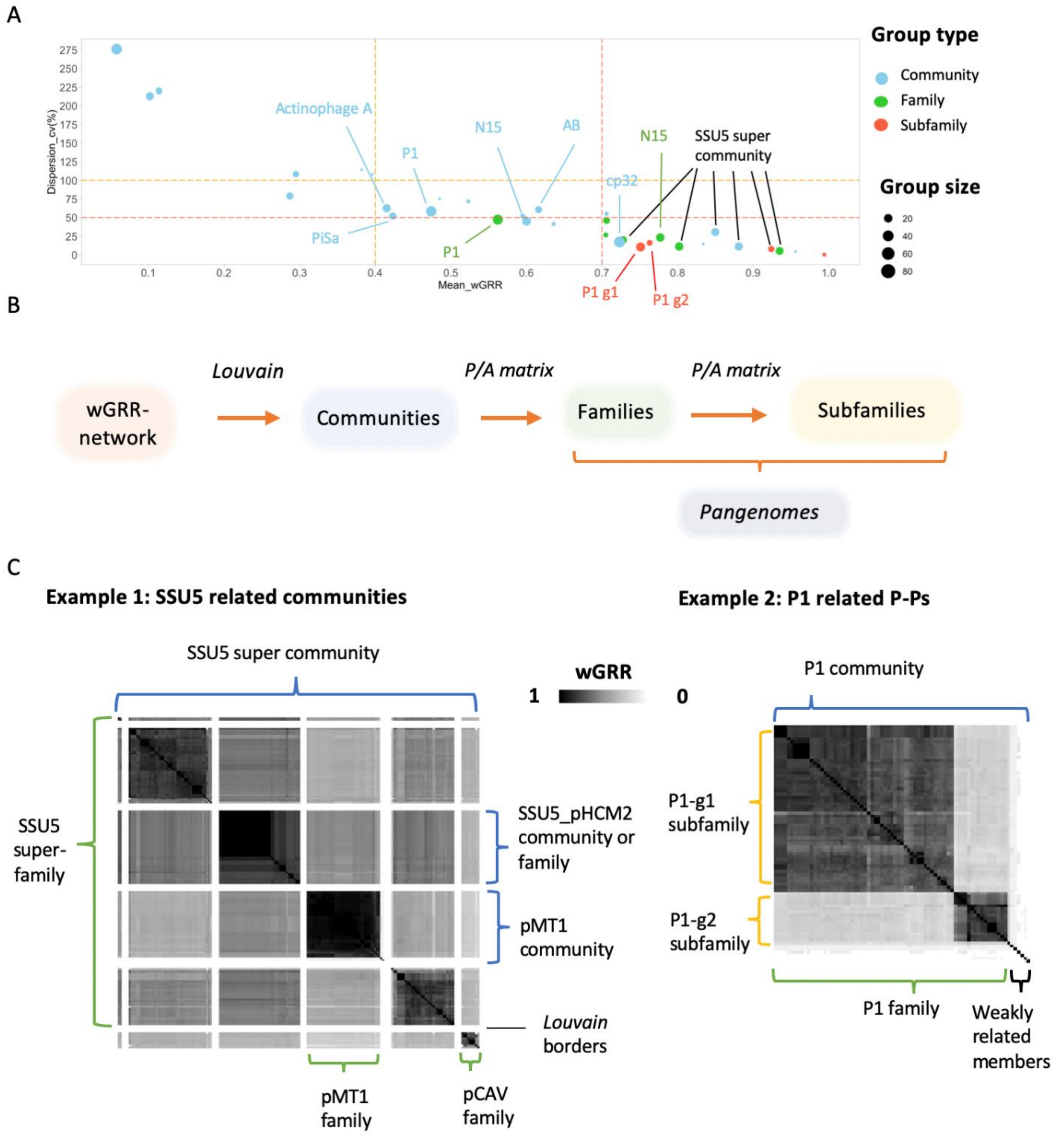

**Figure S4. P-P communities and families.**

**A**. The homogeneity of P-P relatedness within a community was addressed by the coefficient of variation (y-axis) and the mean (x-axis) of all pairwise wGRR scores. Dashed lines separate three areas: (i) highly homogeneous communities ($mean_{wGRR} > 0.7$, coefficient of variation (cv) < 50%), (ii) intermediate communities ($mean_{wGRR} > 0.4$, cv < 100%) and (iii) highly diverse communities. **B**. P-P communities were assigned by the Louvain detection algorithm. P-P Communities were curated into families or subfamilies using the wGRR and the pangenomes. **C**. Examples of P-P (super) communities, (super) families, subfamilies based on the wGRR similarity heatmap.

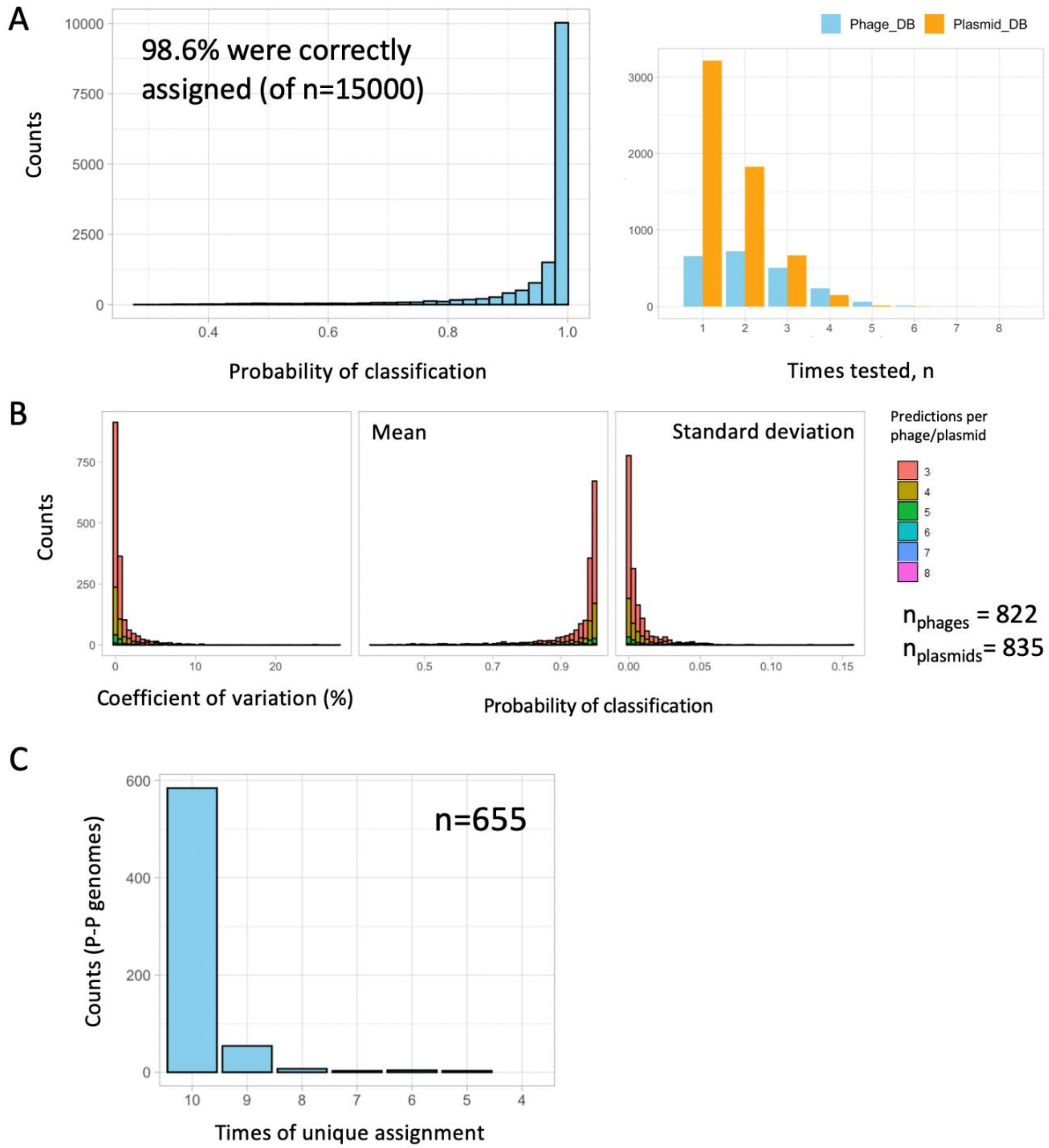

**Figure S5. Prediction of the virus taxonomy of P-Ps using random forest models.**
**A**. 10 random forest models were trained each on 2000 phages (positive cases) and 2000 plasmids (negative cases). The evaluation was done using 10 test datasets each consisting of 1500 randomly chosen replicons (500 phages, 1000 plasmids). For each model, train and test datasets were chosen to be independent. Of the 15000 test cases, 98.6% were assigned correctly. Left panel: distribution of the predictions. Right panel: counts of phage and plasmid classifications due to the random sampling (a taxonomy was assigned to some phages and plasmids several times). **B**. Analysis of elements that were classed by the models at least three times (822 phages and 835 plasmids). Shown are the mean, standard deviation and coefficient of variation (cv) distributions of the probability scores. **C**. The random forest models were used to classify the virus taxonomy of 655 P-Ps. For each P-P a taxonomy was predicted ten times (once per model). In 584 cases the classifications were consistent. For the remaining 71 P-Ps, multiple taxonomies were assigned and therefore only the ones with the highest frequencies were chosen.

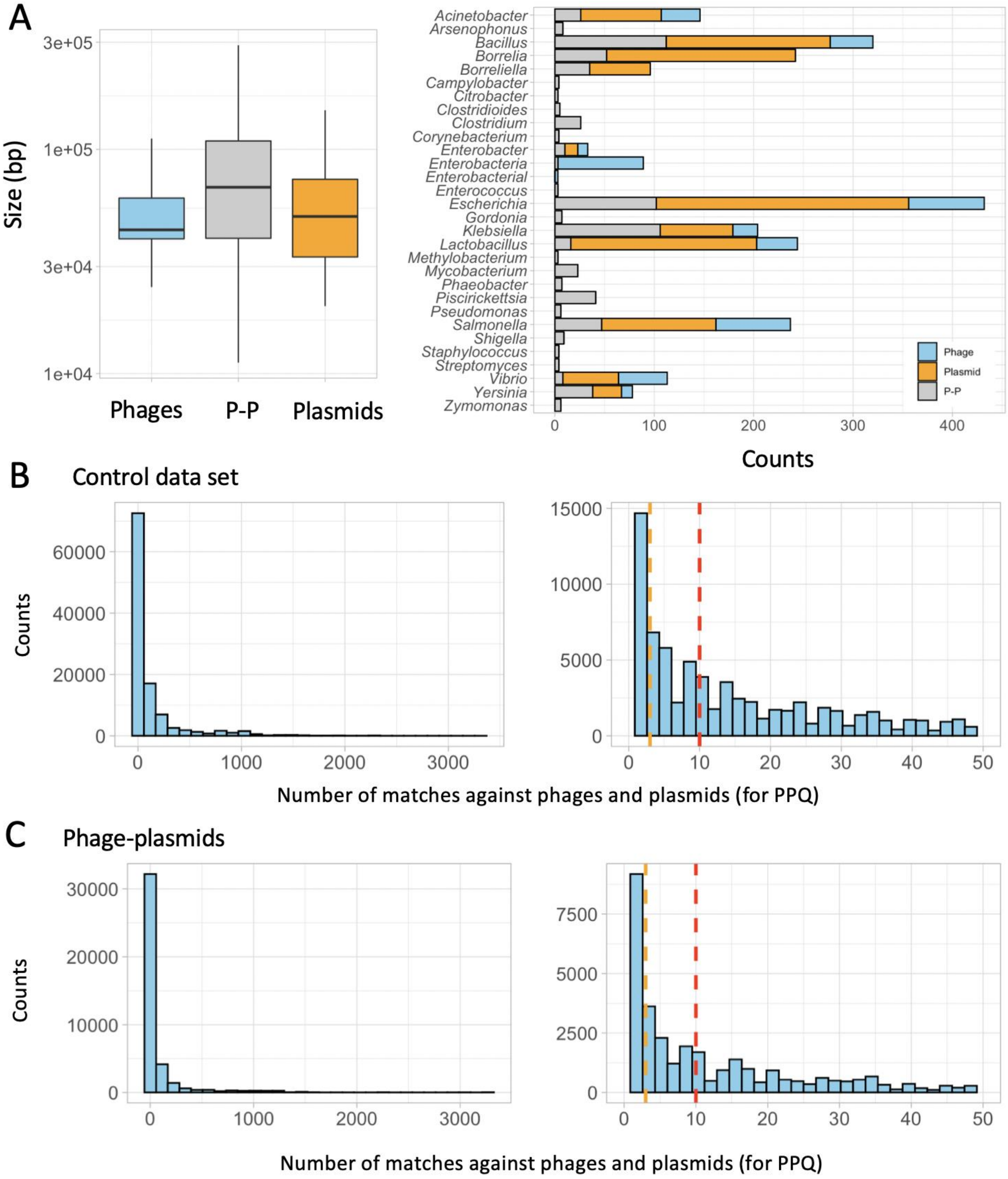

**Figure S6. Evaluation of the phage-plasmid quotient (PPQ).**

**A**. Size (left panel) and host distribution (right panel) of 677 P-Ps, 460 phages and 1226 plasmids (control cases) that were selected (see Methods) to evaluate the PPQ scores. **B and C**. Counts of proteins that match phages or plasmids (for the control and the P-P data set). Left panels show the full range of the hits (from 1 up to >3300 hits) and the right panels zoomed regions (1 up to 50). Dashed lines indicate the thresholds used in the PPQ pangenome graphs for the color intensity (<3, >3 & < 10, >10).

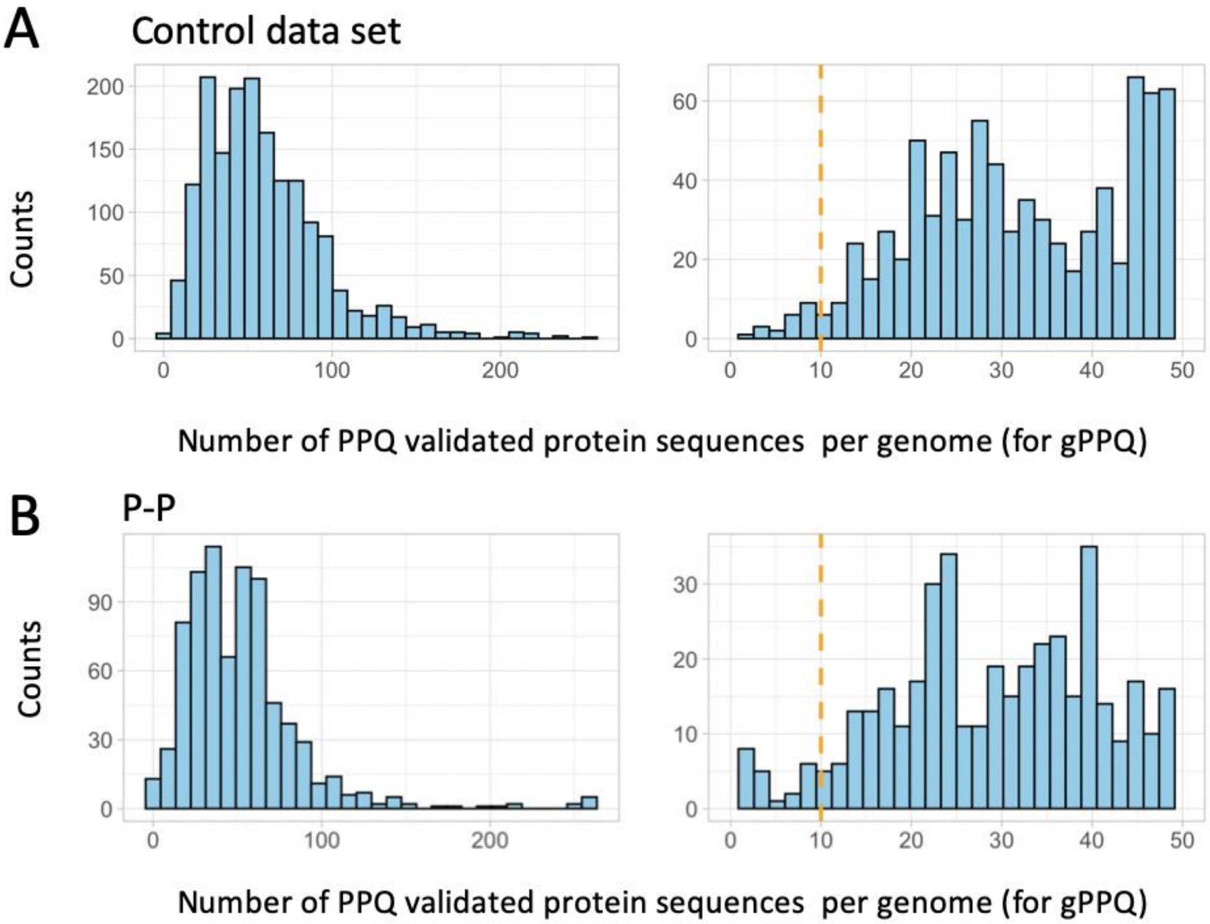

**Figure S7. Computation of the gPPQ is based on replicons with at least ten protein sequences for which a PPQ could be computed.**

**A and B**. Shown are the counts of replicons (control phages and plasmids in A; P-Ps in B) according to the number of proteins used to compute the gPPQ (for details see Methods). Left panels show the full range (1 up to >250 PPQ considered sequences per replicon) and right panels zoomed in region (1 up tp 50 PPQ protein sequences). The gPPQ scores (shown in Fig. 4 CD) were calculated only for replicons that contain at least 10 PPQ validated sequences (orange dashed line).

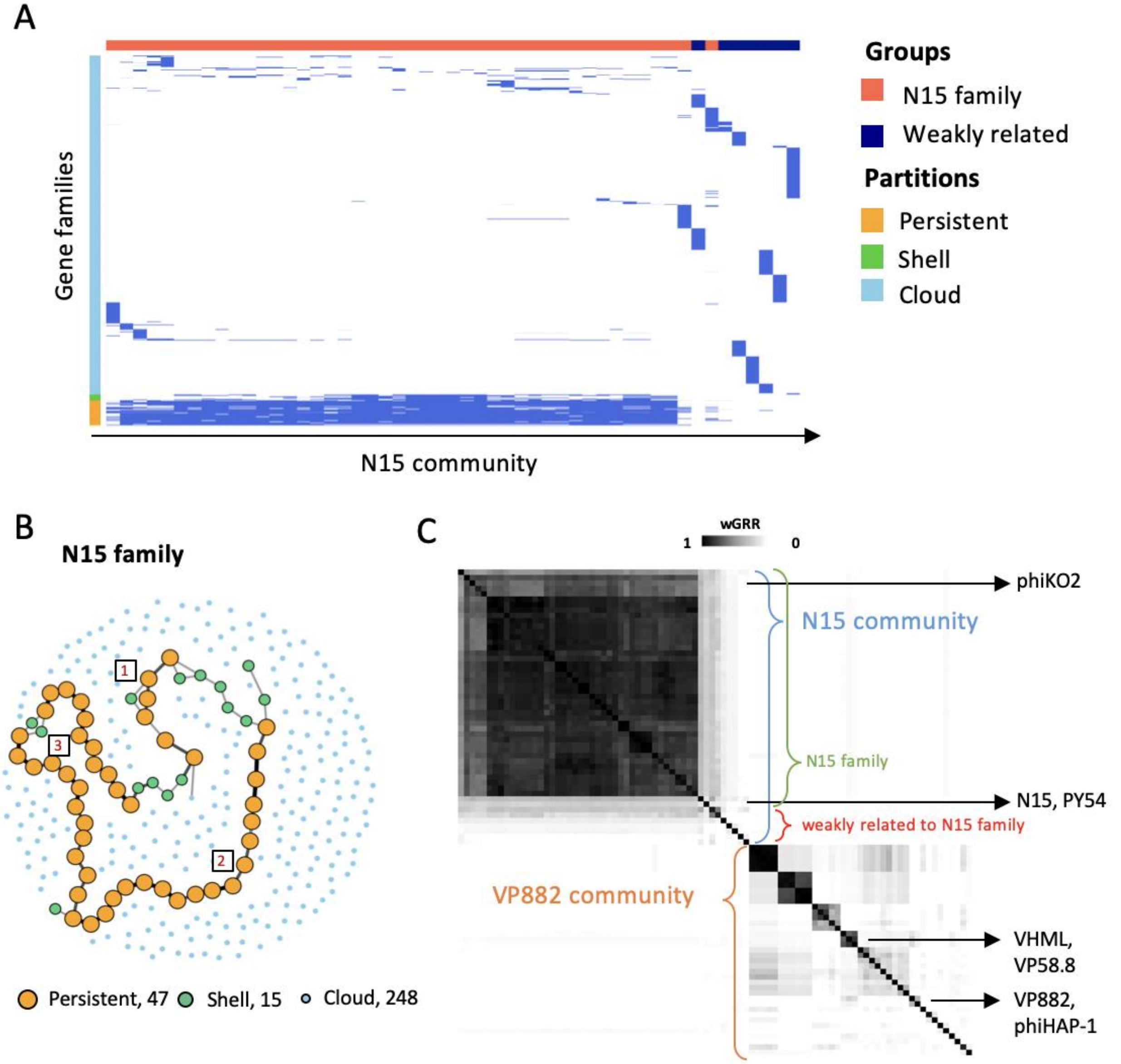

**Figure S8. Pangenome of the N15 family.**

**A**. Presence/absence matrix of the of the genomes of the N15 community classified in persistent, shell and cloud.

**B**. Pangenome graph of the of the N15 family. Nodes are gene families. Size and color indicate the different types of gene families (persistent, shell and cloud). Edges represent neighborhood between the genes. No edge is drawn when the frequency of contiguity is lower than 25%. Grey/thin: moderately co-localized (25 – 50%, 50-90%). Black/thick: in >90% of the genomes. The number of large conserved regions are identified by numbers in boxes. **C**. wGRR relatedness matrix of the N15 (blue) and VP882 community (orange).

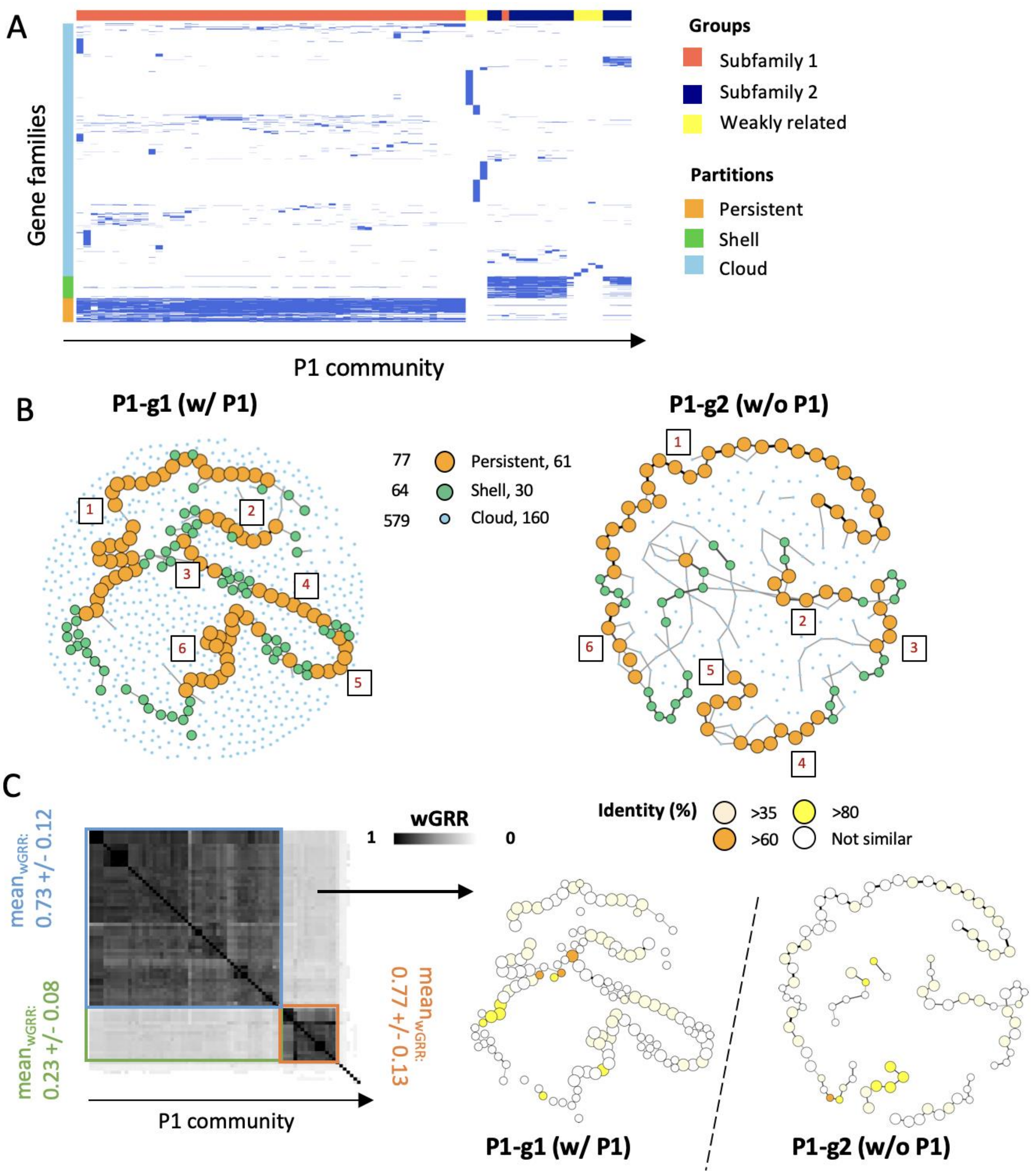

**Figure S9. Curation and comparison of the P1 community.**

**A**. Presence/absence matrix of the of the genomes of the P1 community classified in persistent, shell and cloud (for the subfamilies). **B**. Pangenomes of the two P1 subfamilies. Nodes (colors and size) represent different types of gene families. Edges represent neighborhood between the genes. No edge is drawn when the frequency of contiguity is lower than 15%. Grey/thin: moderately co-localized (15 – 50%, 50-90%). Black/thick: in >90% of the genomes. Large conserved regions are indicated by numbers in boxes. **C**. Left panel: wGRR based heatmap of the P1 community shows the relation between subfamily1 and 2. Right panel: Similarity pangenome graphs of the two subfamilies. Gene families that contain BBH (from one subfamily to the other) are pointed out by red colors (depending on the average protein identity).

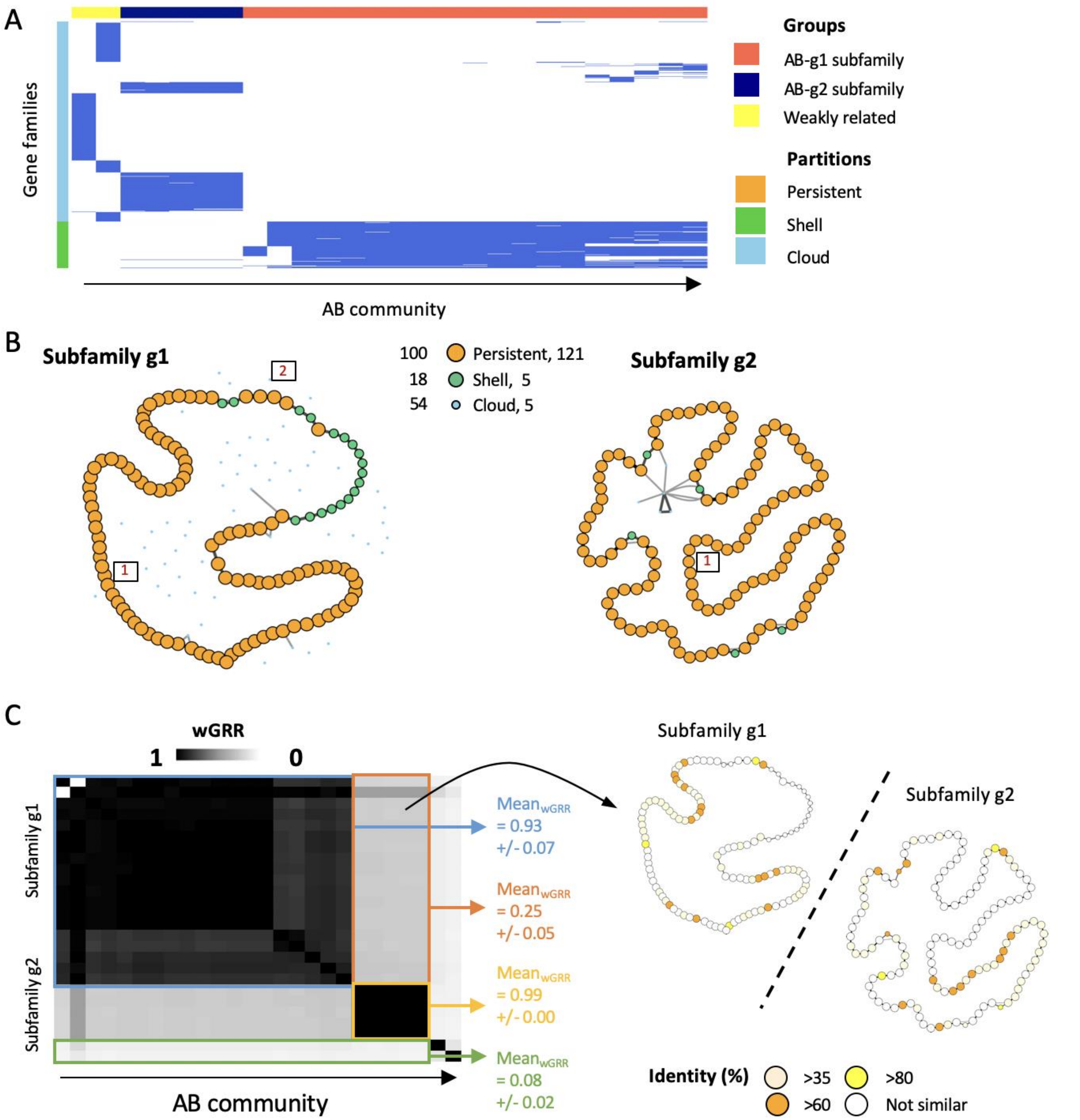

**Figure S10. Pangenome based curation of the AB community.**

**A**. Presence/absence of gene families of the AB community partitioned into persistent, shell and cloud genome. Differentiation of subfamilies were done using a common shell genome (at least 10%). **B**. Pangenome graphs of the two AB subfamilies. Nodes are gene families. Types are indicated by size and color. Edges represent neighborhood between the genes. No edge is drawn when the frequency of contiguity is lower than 15%. Grey/thin: moderately co-localized (15 – 50%, 50-90%). Black/thick: in >90% of the genomes. Conserved large regions are indicated by numbers in boxes. **C**. Left panel: The wGRR similarity within the AB community is shown by the wGRR heatmap. Means and standard deviations of the groups are indicated by different colors. Right panel: Similarity pangenome graphs of the two AB subfamilies. White nodes represent not related and red nodes show related gene families. Red color intensity depends on the average protein identity of the similar gene families.

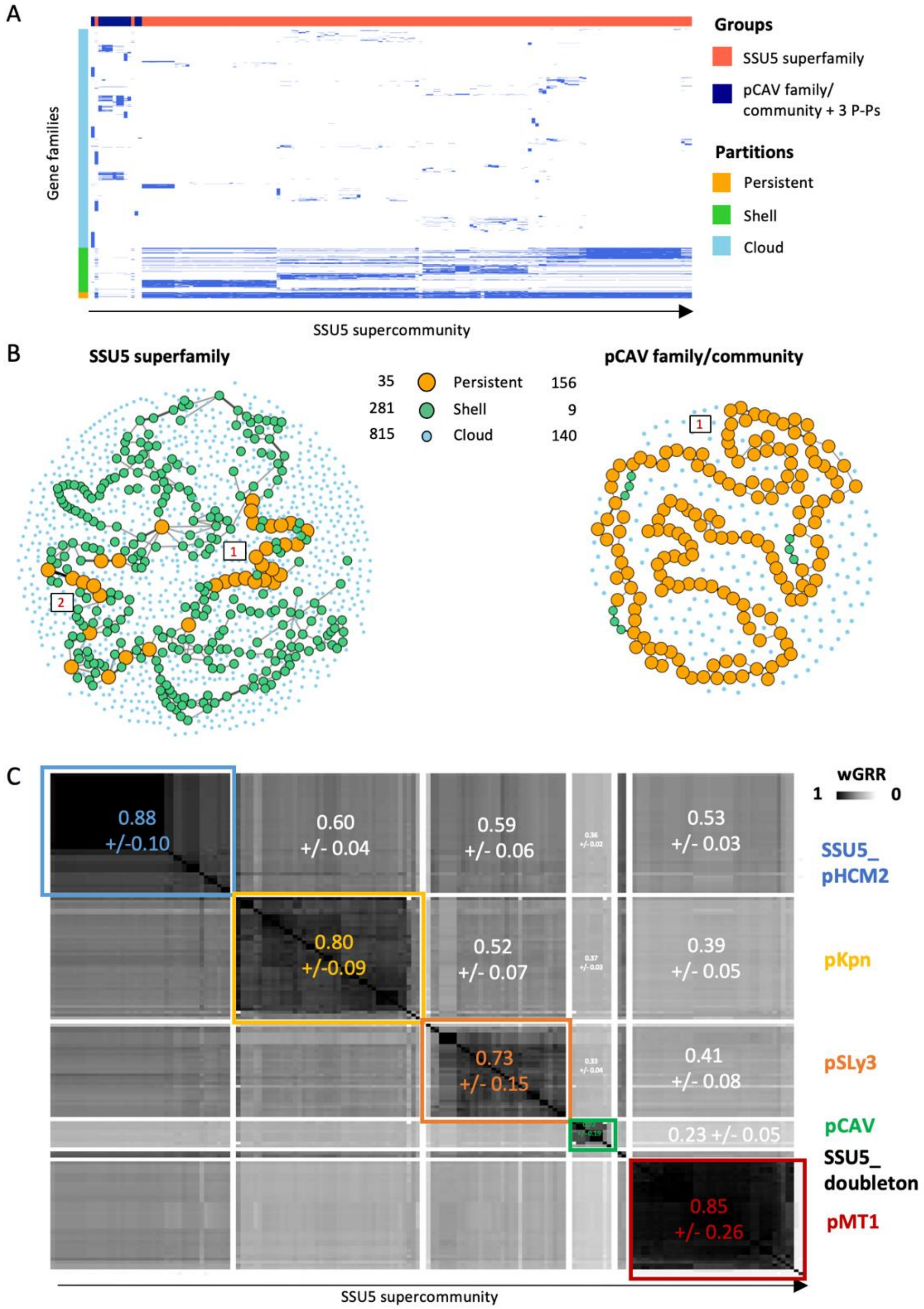

**Figure S11. Comparative analysis of the SSU5 community.**

**A.** Presence/absence matrix of the of the genomes of the SSU5 supercommunity classified in persistent, shell and cloud. Only P-Ps that contain 10% of the persistent genome were assigned to the SSU5 superfamily. **B.** Graphs of the SSU5 super family and pCAV family pangenomes. As described in the figures above, nodes are gene families. Edges represent neighbourhood between the genes. No edge is drawn when the frequency of contiguity is lower than 15%. Grey/thin: moderately co-localized (15 – 50%, 50-90%). Black/thick: in >90% of the genomes. **C.** wGRR similarity heatmap of the SSU5 community. Families are shown in different colors. Mean and standard deviation of and between the communities are indicated by the same color choice.

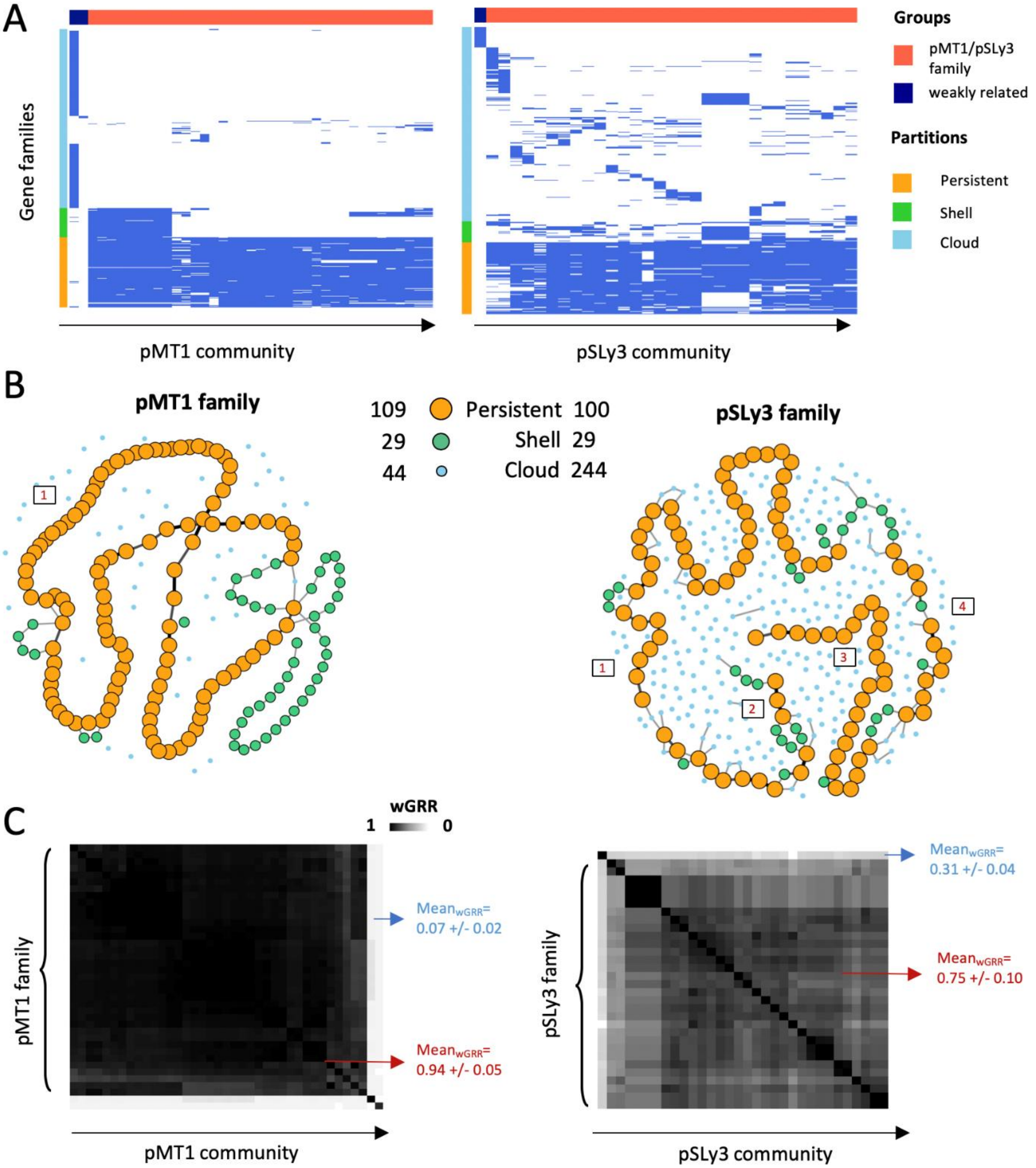

**Figure S12: Curation of the pMT1 and pSLy3 community.**

**A.** Presence/absence matrix of the of the genomes of two communities pMT1 and pSLy3 classified in persistent, shell and cloud (for the subfamilies). **B.** Pangenomes the pMT1 and pSLy3 families. Nodes are gene families, which different types are indicated by colors and size. Edges represent neighborhood between the genes. No edge is drawn when the frequency of contiguity is lower than 15%. Grey/thin: moderately co-localized (15 – 50%, 50-90%). Black/thick: in >90% of the genomes. Number of conserved regions is indicated in boxes. **C.** wGRR heatmaps of the two communities. Mean and standard deviation for the different subgroups of a community are shown in blue/red.

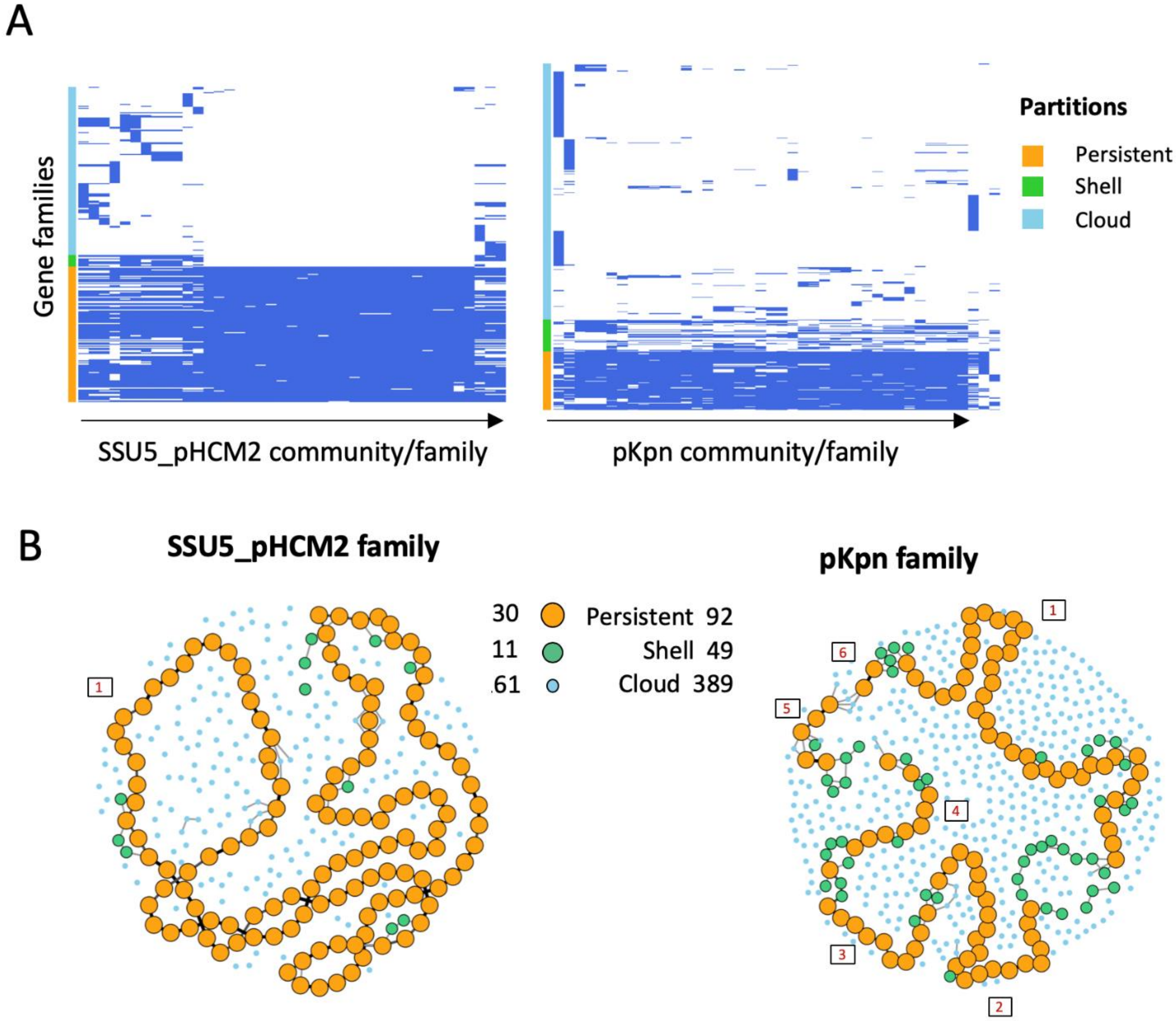

**Figure S13: Pangenomes of the SSU5_pHCM2 and pKpn families.**

**A**. P/A matrixes of the SSU5_pHCM2 and pKpn families computed by PPanGGOLiN (7). A curation was not needed, since all members contain at least 10% of the persistent genome. **B.** Pangenome graphs of the two families. Nodes represent gene families, which different types are distinguished by colors and size. Edges represent neighborhood between the genes. No edge is drawn when the frequency of contiguity is lower than 15%. Grey/thin: moderately co-localized (15 – 50%, 50-90%). Black/thick: in >90% of the genomes. Number of conserved regions was manually assigned and is shown in boxes.

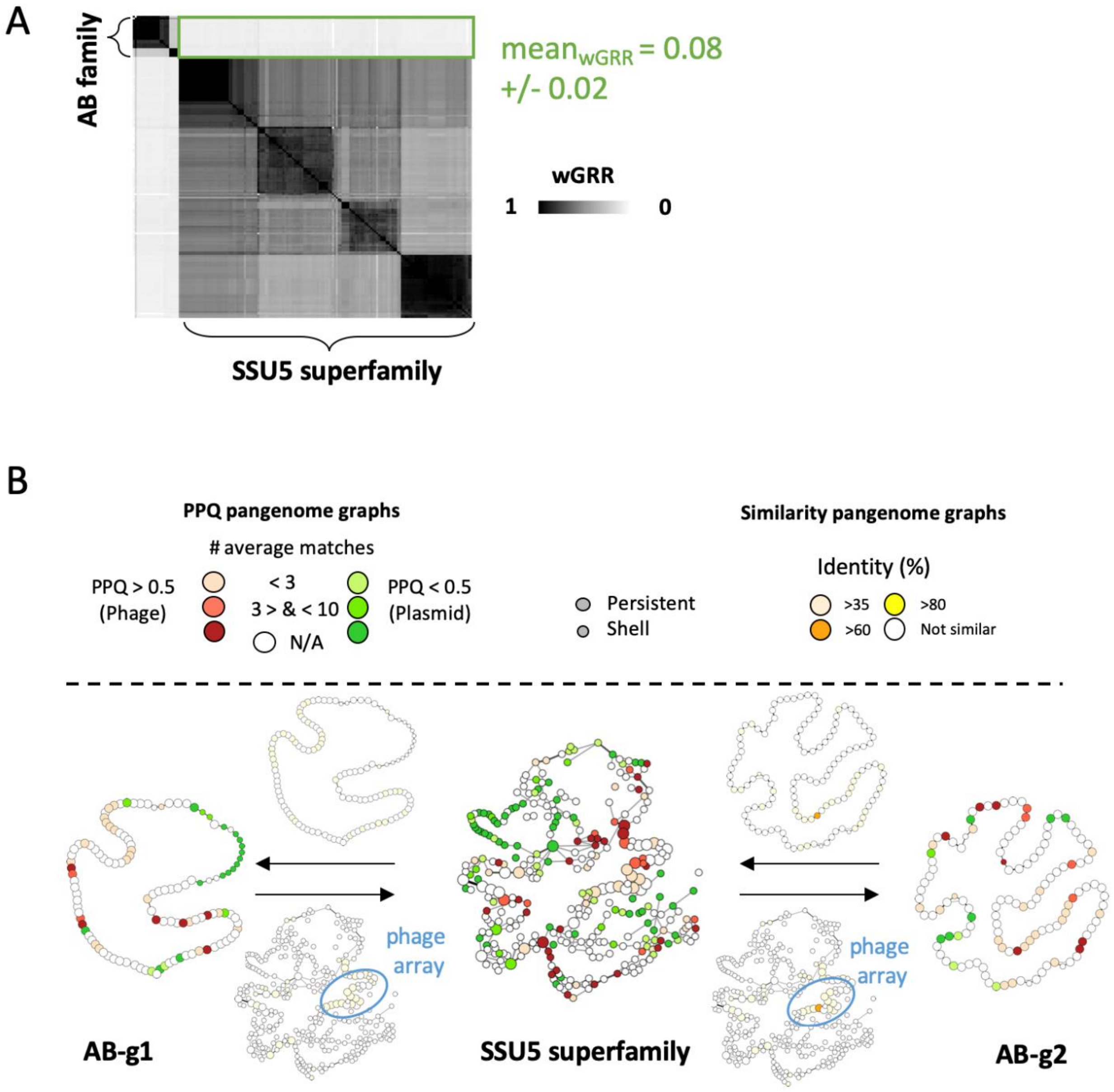

**Figure S14: Comparative analysis of the AB family and SSU5 superfamily.**

**A.** wGRR similarity between the two groups is shown in the wGRR heatmap. **B.** PPQ-Pangenomes graphs of the two (super-) families were compared based on the BBH similarity matrix. In the similarity pangenome graphs (above/under the arrows), gene families of one AB subfamily that contain BBHs to the SSU5 superfamily (above the arrow) are shown in red nodes (vice versa, under the arrows). Red color intensity indicates the average sequence identity.

### N15 family

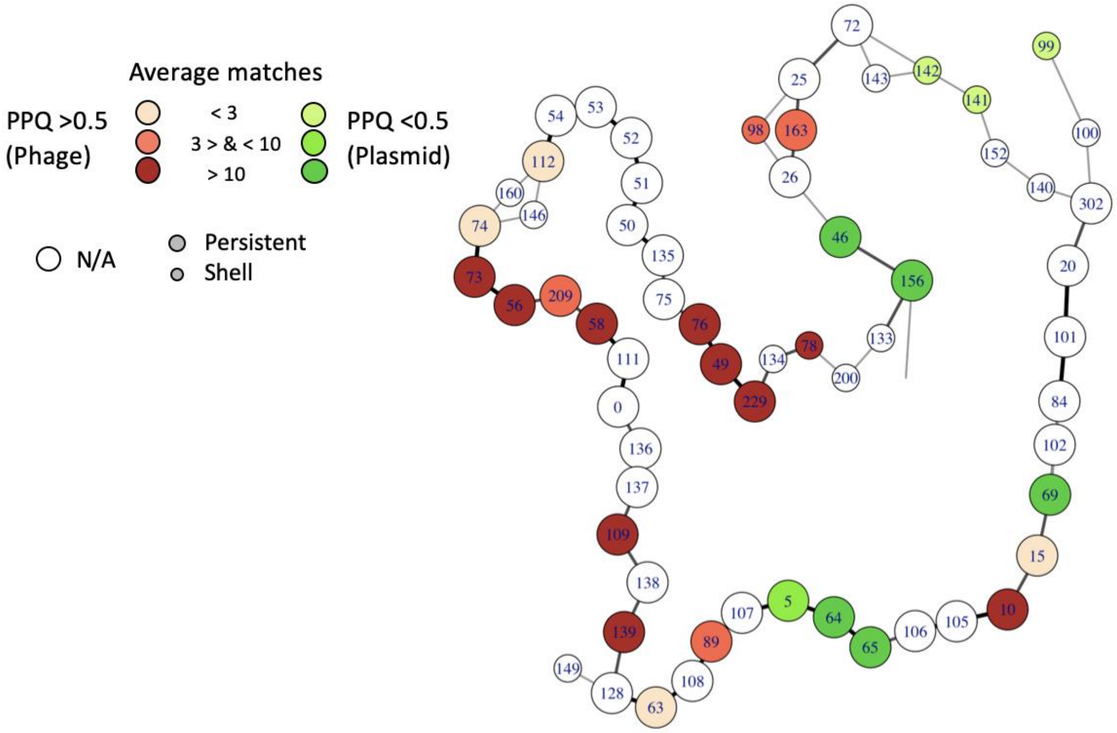

**Figure S15: Indexed pangenome graph of the N15 family.**
Numbers in the nodes show the index of the gene families (suppl. table 9). For details see Figure S8.

### P1 family

#### P1 subfamily 1

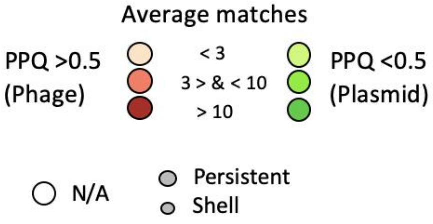

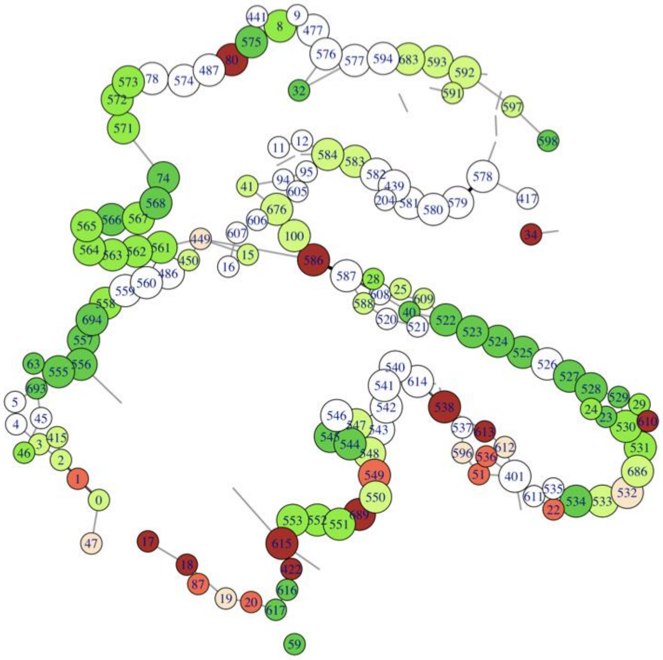

#### P1 subfamily 2

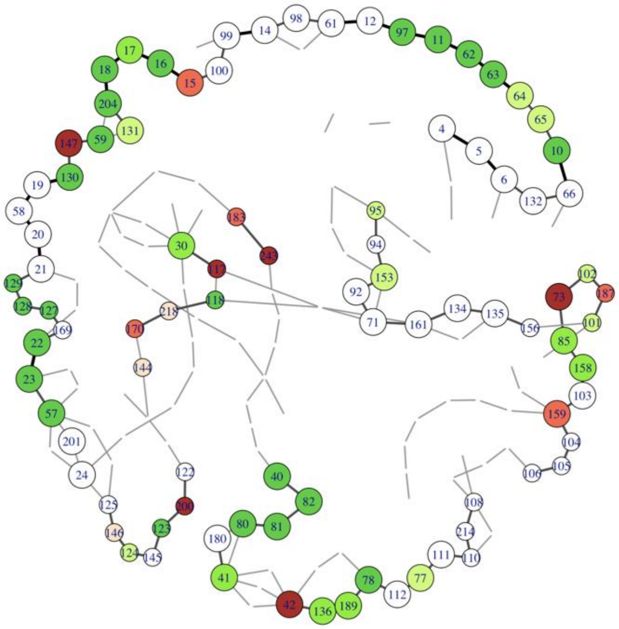

**Figure S16: Indexed pangenome graph of the P1 family.**
For details see Figure S15. Numbers in the nodes show the index of the gene families that are listed in supplementary table 10 and 11.

### AB family

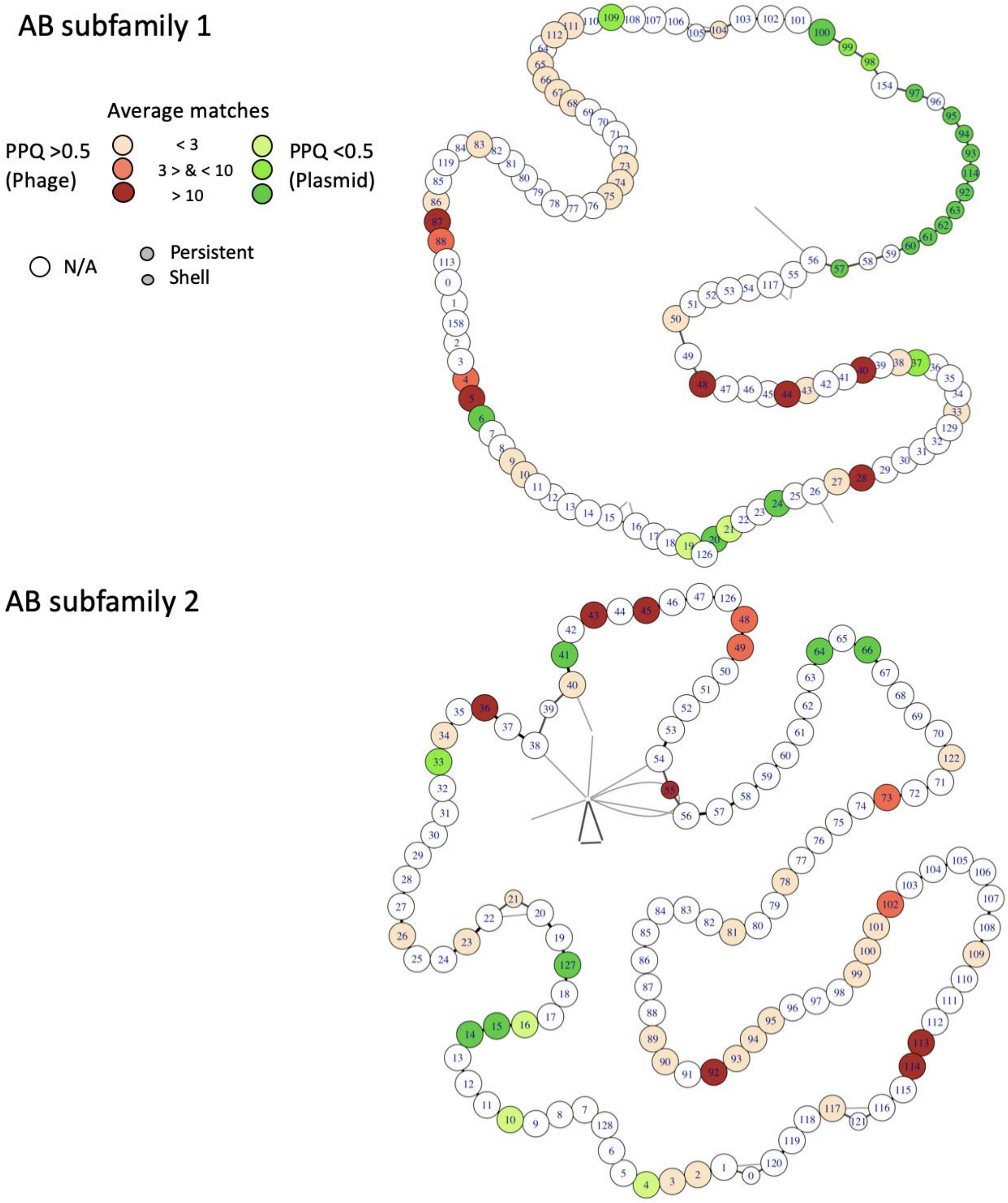

**Figure S17: Indexed pangenome graph of the AB family.**
As Figure S15 but for the AB family. Numbers in the nodes show the index of the gene families that are listed in supplementary table 12 and 13.

### SSU5 superfamily

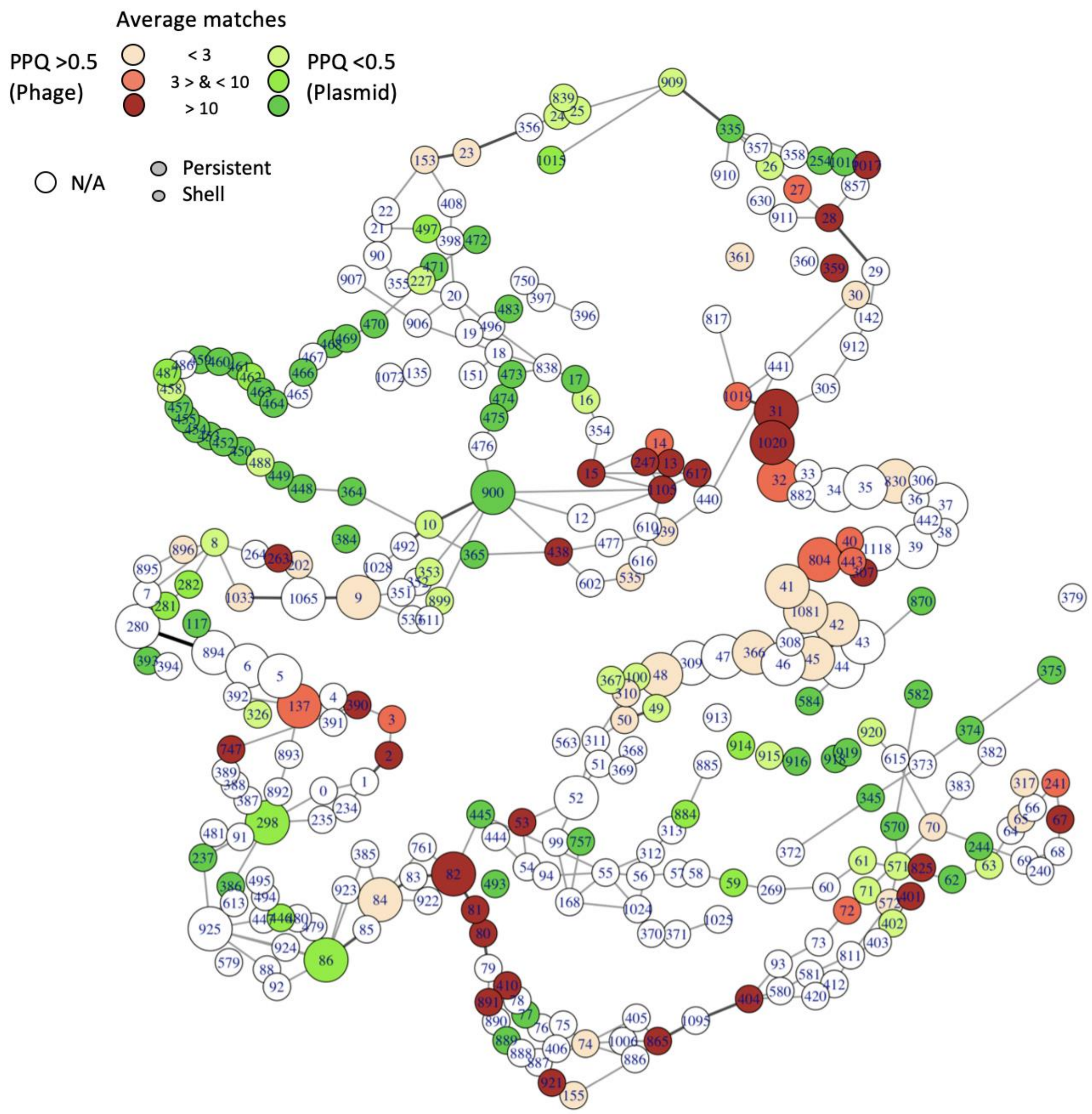

**Figure S18: Indexed pangenome graph of the SSU5 superfamily.**
For details see Figure S15. Numbers in the nodes show the index of the gene families that are listed in supplementary table 18.

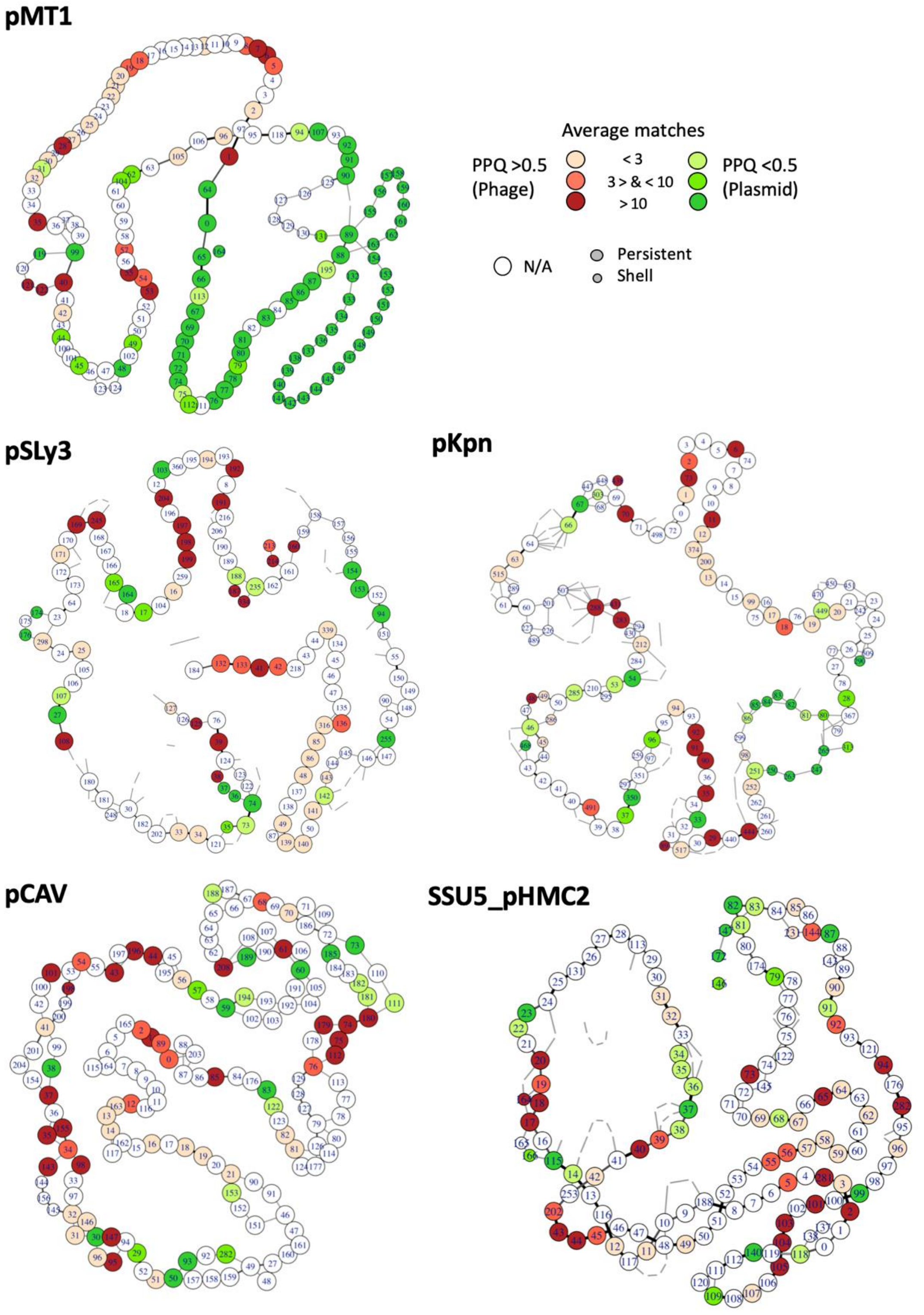

**Figure S19: Indexed pangenome graph of the SSU5-related families.**
As Figure S15 but for the pMT1, pCAV, pSLy3, pKpn and SSU5_pHCM2 families. Numbers in the nodes show the index of the gene families that are listed in supplementary table 14 to 18.

#### REFERENCES

1. Ackermann,H.-W. (2007) 5500 Phages examined in the electron microscope. *Arch Virol*, **152**, 227–243.

2. Grazziotin,A.L., Koonin,E.V. and Kristensen,D.M. (2017) Prokaryotic Virus Orthologous Groups (pVOGs): a resource for comparative genomics and protein family annotation. *Nucleic Acids Res*, **45**, D491–D498.

3. Arndt,D., Grant,J.R., Marcu,A., Sajed,T., Pon,A., Liang,Y. and Wishart,D.S. (2016) PHASTER: a better, faster version of the PHAST phage search tool. *Nucleic Acids Res*, **44**, W16–W21.

4. Robin,X., Turck,N., Hainard,A., Tiberti,N., Lisacek,F., Sanchez,J.-C. and Müller,M. (2011) pROC: an open-source package for R and S+ to analyze and compare ROC curves. *BMC Bioinformatics*, **12**, 77.

5. Christensen,A.P. (2018) NetworkToolbox: Methods and Measures for Brain, Cognitive, and Psychometric Network Analysis in R. *R J.*, **10**, 422–439.

6. Blondel,V.D., Guillaume,J.-L., Lambiotte,R. and Lefebvre,E. (2008) Fast unfolding of communities in large networks. *J. Stat. Mech.*, **2008**, P10008.

7. Gautreau,G., Bazin,A., Gachet,M., Planel,R., Burlot,L., Dubois,M., Perrin,A., Médigue,C., Calteau,A., Cruveiller,S., *et al.* (2020) PPanGGOLiN: Depicting microbial diversity via a partitioned pangenome graph. *PLoS Comput. Biol.*, **16**, e1007732.
